# Avoidance of hydrogen sulfide is modulated by external and internal states in *C. elegans*

**DOI:** 10.1101/2023.10.09.561535

**Authors:** Longjun Pu, Lina Zhao, Jing Wang, Clementine Deleuze, Lars Nilsson, Johan Henriksson, Patrick Laurent, Changchun Chen

## Abstract

Hydrogen sulfide (H_2_S) acts as an energy source, a toxin, and a gasotransmitter across diverse biological contexts. We use the robust locomotory responses of *Caenorhabditis elegans* to high levels of H_2_S to elucidate the molecular mechanisms underlying its acute and adaptive responses. We find that the H_2_S-evoked behavioral response is shaped by multiple environmental factors including oxygen (O_2_) levels and nutritional state, and is modulated by various pathways such as insulin, TGF-β, and HIF-1 signaling, as well as by input from O_2_-sensing neurons. Prolonged exposure to H_2_S activates HIF-1 signaling, leading to the upregulation of stress-responsive genes, including those involved in H_2_S detoxification. This promotes an adaptive state in which locomotory speed is reduced in H_2_S, while responsiveness to other stimuli is preserved. In mutants deficient in HIF-1 signaling, iron storage, and detoxification mechanisms, animals display a robust initial response but rapidly enter a sleep-like behavior characterized by reduced mobility and diminished responsiveness to subsequent sensory stimuli. Furthermore, while acute production of mitochondria-derived reactive O_2_ species (ROS) appears to initiate the avoidance response to H_2_S, persistently high ROS promotes an adaptive state, likely by activating various stress-response pathways, without substantially compromising cellular H_2_S detoxification capacity. Taken together, our study provides comprehensive molecular insights into the mechanisms through which *C. elegans* modulates and adapts its response to H_2_S exposure.

## Introduction

During the late Proterozoic era, the rise in oxygen (O_2_) levels eliminated hydrogen sulfide (H_2_S) from most habitats. Yet, low O_2_ levels, along with high concentrations of hydrogen sulfide (H_2_S) and carbon dioxide (CO_2_), can persist in enclosed environments where bacteria actively decompose organic matter (Olson & Straub, 2016). H_2_S can easily permeate biological membranes and interfere with various cellular processes. One of the most detrimental effects of H_2_S is the disruption of cellular respiration by remodeling the mitochondrial electron transport chain and inhibiting cytochrome *c* oxidase (COX) (Cooper & Brown, 2008; Khan et al., 1990; Nicholls & Kim, 1982; Romanelli-Cedrez, Vairoletti, & Salinas, 2024).

The detection and response to chemical signals are crucial for various organisms to interact effectively with their environment. Animals living in H_2_S-rich environments have evolved mechanisms to either avoid or adapt to these conditions, conferring them survival advantages. Species that encounter periodic increases in H_2_S often exhibit avoidance behaviors. For example, benthic species undertake daily vertical migrations between sulfidic and non-sulfidic waters (Abel, Koenig, & Davis, 1987; Salvanes, Utne-Palm, Currie, & Braithwaite, 2011). Certain metazoan species have seized the ecological opportunities presented by H_2_S-rich environments through the development of cellular adaptations. Livebearing fish species, for instance, have evolved COX subunits that are more resistant to H_2_S and more efficient at H_2_S detoxification (Kelley et al., 2016; Pfenninger et al., 2014).

Free-living nematodes feed on bacteria that thrive in decaying organic matter, such as compost. This complex and dynamic environment, where species like *C. elegans* are commonly found, may exhibit significant fluctuations in H_2_S levels over short distances (Adams, Farwell, Pack, & Bamesberger, 1979; Budde & Roth, 2011; Morra & Dick, 1991; Patange, Breen, Arsuffi, & Ruvkun, 2025; Rodriguez-Kabana, Jordan, & Hollis, 1965; Romanelli-Cedrez et al., 2024). Evidence suggests that *C. elegans* has evolved tolerance to prolonged H_2_S exposure by reprogramming gene expression through H_2_S-induced stabilization of the hypoxia-inducible factor HIF-1 (Budde & Roth, 2010, 2011; Horsman, Heinis, & Miller, 2019; Ma, Vozdek, Bhatla, & Horvitz, 2012; Miller, Budde, & Roth, 2011; Powell-Coffman, 2010; Topalidou & Miller, 2017), and by developing an H_2_S-resistant electron transport chain (Romanelli-Cedrez et al., 2024). In particular, low concentrations of H_2_S are not only well tolerated but can also be beneficial. For instance, exposure to 50 ppm H_2_S has been shown to extend the lifespan and enhance thermotolerance (Fawcett, Hoyt, Johnson, & Miller, 2015; Miller & Roth, 2007). Moreover, endogenous H_2_S production in *C. elegans* modulates its interactions with actinobacteria (Patange et al., 2025). However, exposure to concentrations exceeding 150 ppm proved lethal (Budde & Roth, 2010; Horsman et al., 2019).

In addition to long-term adaptive mechanisms, *C. elegans* exhibits an acute behavioral response to H_2_S exposure (Budde & Roth, 2011). In this study, we investigated whether and how the nematode *C. elegans* acutely avoids environments rich in H_2_S, how it adapts at the molecular and physiological levels to H_2_S exposure, and how these adaptations alter the avoidance response.

## Results

### *C. elegans* increases locomotory activity in response to acute H_2_S exposure

In response to aversive cues, *C. elegans* initiates a pirouette followed by an increase of its locomotory speed to escape the noxious stimuli. When acutely exposed to 150 ppm H_2_S, the laboratory N2 strain exhibited an increased locomotory speed and enhanced turning behavior, while generally decreasing the frequency of reversals, as a strategy to escape the noxious stimuli (***Figure 1A–C, Figure 1—Video 1***). The maximum speed was reached after 6–8 minutes in H_2_S and returned to the baseline over a period of approximately 1 hour (***Figure 1D***). The reduced locomotory speed after 30 minutes in 150 ppm H_2_S was reversible when these animals were immediately challenged with another stimulus, such as near-UV light (***Figure 1E***), suggesting that neuromuscular function remains preserved. The magnitude and dynamics of the H_2_S-induced avoidance responses were concentration-dependent. Under our assay conditions, 50 ppm H_2_S did not elicit an acute speed increase, and responses to 75 ppm were variable. In contrast, exposure to 150 ppm H_2_S consistently triggered a robust and reproducible escape response (***Figure 1F***), consistent with earlier reports that H_2_S at 50 ppm exerts beneficial effects, while H_2_S at 150 ppm is toxic to *C. elegans* (Budde & Roth, 2010; Fawcett et al., 2015; Horsman et al., 2019; Miller & Roth, 2007). To investigate whether low levels of H_2_S might be preferable or even attractive to the animals, we exposed wild type animals to a linear gradient of H_2_S (***Figure 1G***; See methods for details) (Bretscher, Busch, & de Bono, 2008; Chang, Chronis, Karow, Marletta, & Bargmann, 2006; Gray et al., 2004). Animals were allowed to move freely on a thin layer of bacteria within a microfluidic chamber containing an H_2_S gradient ranging from 150 ppm at one end to 0 ppm at the other. As expected, *C. elegans* robustly avoids 150 ppm H_2_S but does not accumulate at the extreme end of 0 ppm ***(Figure 1G, H).*** The highest proportion of animals was found in the region with approximately 40 ppm H_2_S. These data suggest that H_2_S acts as a potent repellent for *C. elegans* at high concentrations but as an attractant at low levels.

**Figure 1.**
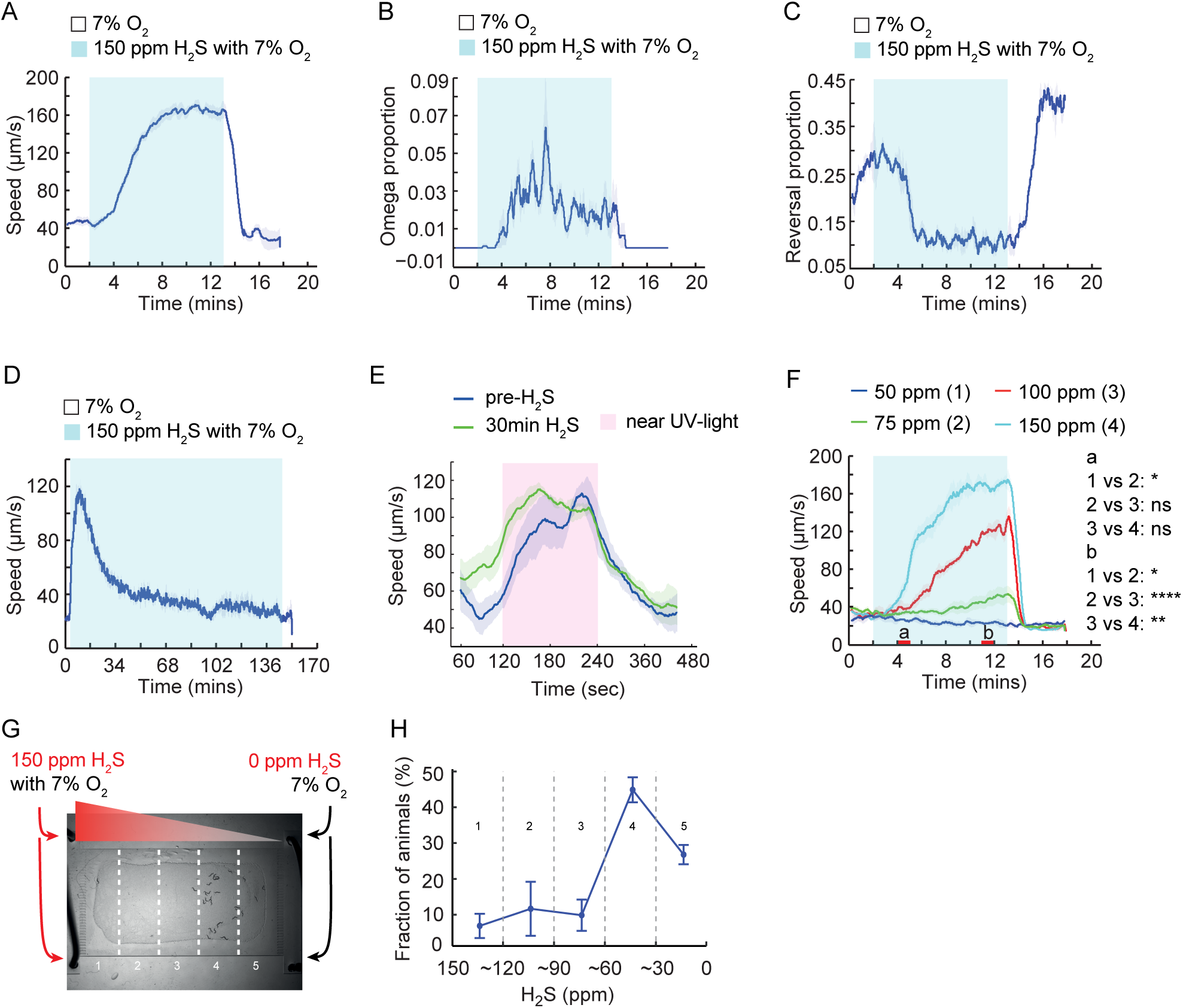
Acute locomotory responses of *C*. *elegans* to hydrogen sulfide. (**A**) Changes in the locomotory speed of WT animals (N2 laboratory strain) evoked by a switch from 7% O_2_ to 150 ppm H_2_S balanced with 7% O_2_, followed by a return to 7% O_2_. (**B**) Proportion of WT animals undergoing reorientation movements (omega turns) during a switch from 7% O_2_ to 150 ppm H_2_S balanced with 7% O_2_, followed by a return to 7% O_2_. (**C**) Fraction of WT animals doing backward locomotion (reversals) during a switch from 7% O_2_ to 150 ppm H_2_S balanced with 7% O_2_, followed by a return to 7% O_2_. (**D**) Locomotory speed of WT was recorded for 2 minutes at 7% O_2_, followed by 148 minutes at 150 ppm H_2_S balanced with 7% O_2_, and then at 7% O_2_. (**E**) Locomotory speed of WT animals in response to near-UV light (435 nm, 0.7 mW/mm^2^) exposure, before and directly after 30-minute preincubation in 150 ppm H_2_S balanced with 7% O_2_. Locomotory activity was recorded for 2 minutes before, during and after UV light exposure. (**F**) Locomotory speed of WT animals to different concentrations of H_2_S balanced with 7% O_2_. Red bars on the x-axis represent two intervals (4–5 minutes and 11–12 minutes, labeled a and b, respectively) used for statistical analysis. **** = *p* < 0.0001, ** = *p* < 0.01, * = *p* < 0.05, ns = not significant, Mann–Whitney U test. (**G**) Distribution of WT animals in a microfluidic device after 25 minutes of aerotaxis. The gas inputs and the five chamber sections used for scoring are indicated. (**H**) Distribution of WT animals in a gradient of H_2_S (150 ppm to 0 ppm). The bins correspond to different sections of the microfluidic chamber described in (**G**). N= 4 aerotaxis assays.

### Candidate gene screen identifies signaling pathways required for H_2_S-evoked locomotion

To investigate the molecular mechanisms underlying the H_2_S-evoked locomotory response in *C. elegans*, we conducted a candidate gene screen for mutants that failed to increase their locomotory speed upon exposure to an acute rise of H_2_S level to 150 ppm. Our analysis prioritized genes that are either known to be, or potentially involved in, sensory responses to gaseous stimuli in both *C. elegans* and mammals (***Supplementary file 1***). We began by examining mutants affecting globins, guanylate cyclases, and cyclic nucleotide-gated (CNG) channels, which are involved in O_2_ or CO_2_ sensing in *C. elegans*, as well as wild isolates and other previously characterized mutants with defective responses to gas stimuli (***Supplementary file 1***) (Bretscher et al., 2008; Gray et al., 2004; Hallem & Sternberg, 2008; Persson et al., 2009). Given the essential role of K^+^ channels in acute hypoxia sensing in mammals, we also screened all potassium (K^+^) channel mutants (Gao, Gonzalez-Rodriguez, Ortega-Saenz, & Lopez-Barneo, 2017; Weir, Lopez-Barneo, Buckler, & Archer, 2005). To explore the neuronal components involved in H_2_S-evoked avoidance, we assayed mutants with defects in ciliogenesis or cilia-mediated signaling, as well as animals deficient in neurotransmission involving classical neurotransmitters, neuropeptides, or biogenic amines. Finally, considering that mitochondrial function is central to H_2_S detoxification, we included mitochondrial electron transport chain (ETC) mutants and key factors implicated in H_2_S and superoxide clearance. Together, these functionally diverse categories allowed us to survey a wide spectrum of potential mechanisms contributing to the behavioral response to H_2_S.

In the screen, a collection of mutants displayed reduced speed response, whereas only a few exhibited a lack of omega-turn and/or reversal responses to H_2_S (***Supplementary file 1***). The reversal inhibition upon H_2_S exposure and its rebound following H_2_S withdrawal appeared to vary substantially across experiments for N2, suggesting that differences in population density, food availability, and animals’ physiological state prior to the assay may matter. Nevertheless, our screen revealed that signaling pathways, including cGMP, insulin and TGF-β signaling, contributed to H_2_S-evoked avoidance (***Supplementary file 1***). A naturally occurring variant of the neuropeptide receptor gene *npr-1*, previously implicated in the modulation of O_2_ and CO_2_ responses, also affected locomotory speed in H_2_S (***Supplementary file 1***). In addition, mitochondrial components, including those involved in respiration and H_2_S detoxification, played critical roles in triggering and maintaining escape behavior in H_2_S. In contrast, we did not find evidence that globins, K^+^ channels, or biogenic amine signaling contributed significantly to the behavioral avoidance to H_2_S (***Supplementary file 1***).

### cGMP signaling in ASJ cilia contributes to H_2_S avoidance

One group of mutants exhibiting reduced locomotory speed in H_2_S had defects in ciliogenesis (***Figure 2A–C***). This group included *daf-19* mutants lacking cilia, *dyf-3* mutants with reduced cilia length, and *dyf-7* mutants in which ciliogenesis occurs but the cilia are not anchored to their sensilla and are not exposed to the external environment (Heiman & Shaham, 2009; Murayama, Toh, Ohshima, & Koga, 2005; Starich et al., 1995; Swoboda, Adler, & Thomas, 2000). These findings suggest that ciliogenesis and exposure of cilia to the external environment are essential for animals to escape H_2_S exposure. In addition, these mutants exhibited consistently high reversal rate, likely associated with their reduced locomotory speed (***Figure 2—figure supplement 1A–F***). Cilia-mediated sensory responses often involve the activation of guanylate cyclases and the opening of cGMP-gated channels (Ferkey, Sengupta, & L’Etoile, 2021). We observed significantly attenuated locomotory speed in response to H_2_S in *tax-4* or *tax-2* mutants, which lack key subunits of cilia-enriched cGMP-gated channels (***Figure 2D, E***). In a comprehensive survey of all guanylate cyclase mutants, the *daf-11* strain emerged as the only mutant showing speed response defects in the avoidance of H_2_S (***Figure 2D, supplementary file 1***). Similar to the cilia mutants, animals deficient in *daf-11* had low speed, attenuated omega turns, and constantly high reversal rate (***Figure 2D, Figure 2—figure supplement 1G, H***). Selective expression of *daf-11* in ASJ neurons, but not in other neurons, partially restored the speed response to acute H_2_S (***Figures 2F, Figure 2—figure supplement 2A–D***). In addition, the H_2_S response defect of *tax-4* mutants was also rescued by selective expression of its cDNA in ASJ neurons (***Figure 2G***), suggesting that the receptor guanylate cyclase DAF-11 and the CNG channel TAX-4 act at least partially in ASJ neurons to promote acute avoidance to H_2_S. To validate the role of ASJ neurons in H_2_S responses, we blocked neurotransmission from ASJ using the catalytic domain of *Tetanus Toxin (TeTx*), which impairs neurosecretion by specifically cleaving synaptobrevin (Schiavo et al., 1992). Cell-specific expression of *TeTx* in ASJ significantly inhibited the speed response to H_2_S ***(Figure 2H)***. Surprisingly, using the Ca^2+^ sensor GCaMP6s to monitor a direct response of ASJ neurons to H_2_S, we observed no H_2_S-induced Ca^2+^ transients under any of the tested conditions, whereas CO_2_-evoked Ca^2+^ increases were readily detected ***(Figure 2I, Figure 2—figure supplement 2E).*** These findings suggest that although ASJ neuronal activity and neurosecretion contribute to the H_2_S responses, ASJ neurons are unlikely to play a role in directly detecting H_2_S.

**Figure 2.**
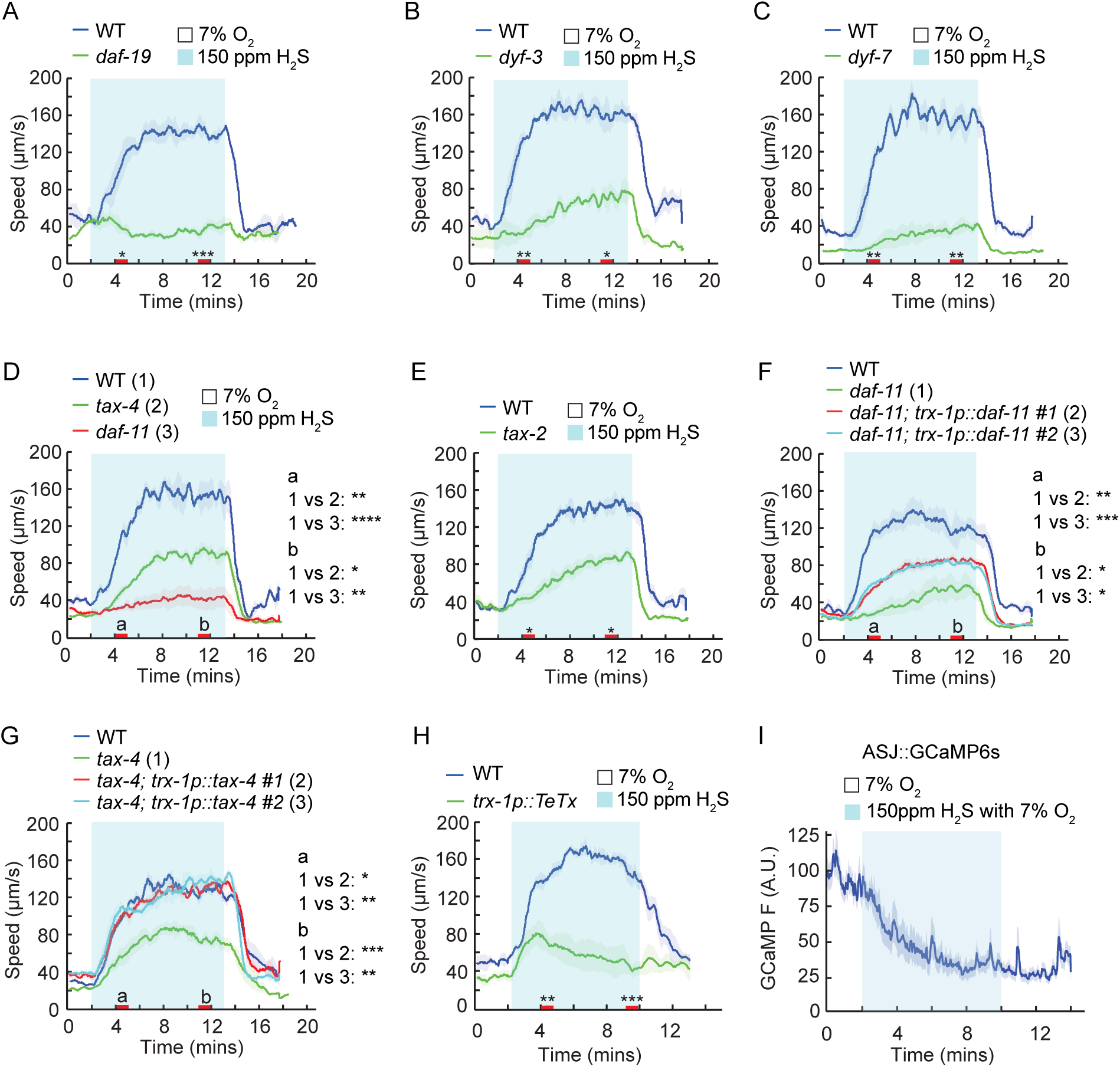
Neurosecretion from ASJ neurons contributes to H_2_S avoidance. (**A–H**) Locomotory speed responses to a switch from 7% O_2_ to 150 ppm H_2_S balanced with 7% O_2_ for animals of the indicated genotype: WT and *daf-19(m86)* (**A**); WT and *dyf-3(m185)* (**B**); WT and *dyf-7(m539)* (**C**); WT, *daf-11(m47)* and *tax-4(p678)* (**D**); WT and *tax-2(p691)* (**E**); WT, *daf-11(m47)* and transgenic *daf-11(m47)* expressing *daf-11* genomic DNA in ASJ neurons (two independent lines) (**F**); WT, *tax-4(p678)* and transgenic *tax-4(p678)* expressing *tax-4* cDNA in ASJ neurons (two independent lines) (**G**); WT and transgenic WT expressing the catalytic domain of the *tetanus* toxin (TeTx) in ASJ neurons (**H**). For the comparison of acute locomotory speed responses between strains, red bars on the x-axis represent two intervals (4– 5 minutes and 11–12 minutes, labeled a and b, respectively, in panels D, F, and G) used for statistical analysis. **** = *p* < 0.0001, *** = *p* < 0.001, ** = *p* < 0.01, * = *p* < 0.05, ns = not significant, Mann–Whitney U test. (**I**) Calcium transients evoked in ASJ neurons of WT animals in response to a switch from 7% O_2_ to 150 ppm H_2_S balanced with 7% O_2_ for animals (N=42). The slow decline in GCaMP6s fluorescence is due to photobleaching.

### Starvation modulates H_2_S avoidance

In the candidate gene screen, another set of mutants with reduced speed responses to H_2_S had previously been shown to be defective in response to CO_2_ (Hallem & Sternberg, 2008). Specifically, the H_2_S-evoked speed response was reduced in mutants deficient in nutrient-sensitive signaling, including the insulin receptor DAF-2, the TGF-β ligand DAF-7, and the nuclear hormone receptor NHR-49 (***Figure 3A–C, Figure 3—figure supplement 1A***). Similarly, mutants lacking other CO_2_ response modulators, such as the calcineurin subunits TAX-6 and CNB-1, also failed to respond to H_2_S (***Figure 3—figure supplement 1B, C***). In addition to the abolished speed response, we also observed that the H_2_S-evoked omega-turn response was nearly abolished in *daf-2* mutants, and high proportion of animals exhibited persistently high reversal rate regardless of H_2_S stimulation (***Figure 3—figure supplement 1D–F***). The H_2_S avoidance defects observed in *daf-2* and *daf-7* single mutants were fully suppressed in *daf-2; daf-16* and *daf-7; daf-3* double mutants ***(Figure 3A, B).*** Therefore, it is likely that insulin and TGF-β signaling modulate H_2_S responses by regulating the expression of relevant genes via DAF-16 and DAF-3 transcription factors, respectively. Given that H_2_S-evoked speed response was modulated by the nutrient-sensitive pathways, we further explored whether the locomotory response to H_2_S was sensitive to the nutrient state. Similar to the response to CO_2_ (Bretscher et al., 2008; Hallem & Sternberg, 2008), we observed that a 24-hour period of starvation suppressed the speed response to H_2_S ***(Figure 3D).*** However, the CO_2_ sensor GCY-9 was dispensable for H_2_S-evoked avoidance ***(Figure 3—figure supplement 1G–I).*** These findings suggest that while acute avoidance of CO_2_ and H_2_S share common modulatory pathways, the responses are initiated through distinct mechanisms.

**Figure 3.**
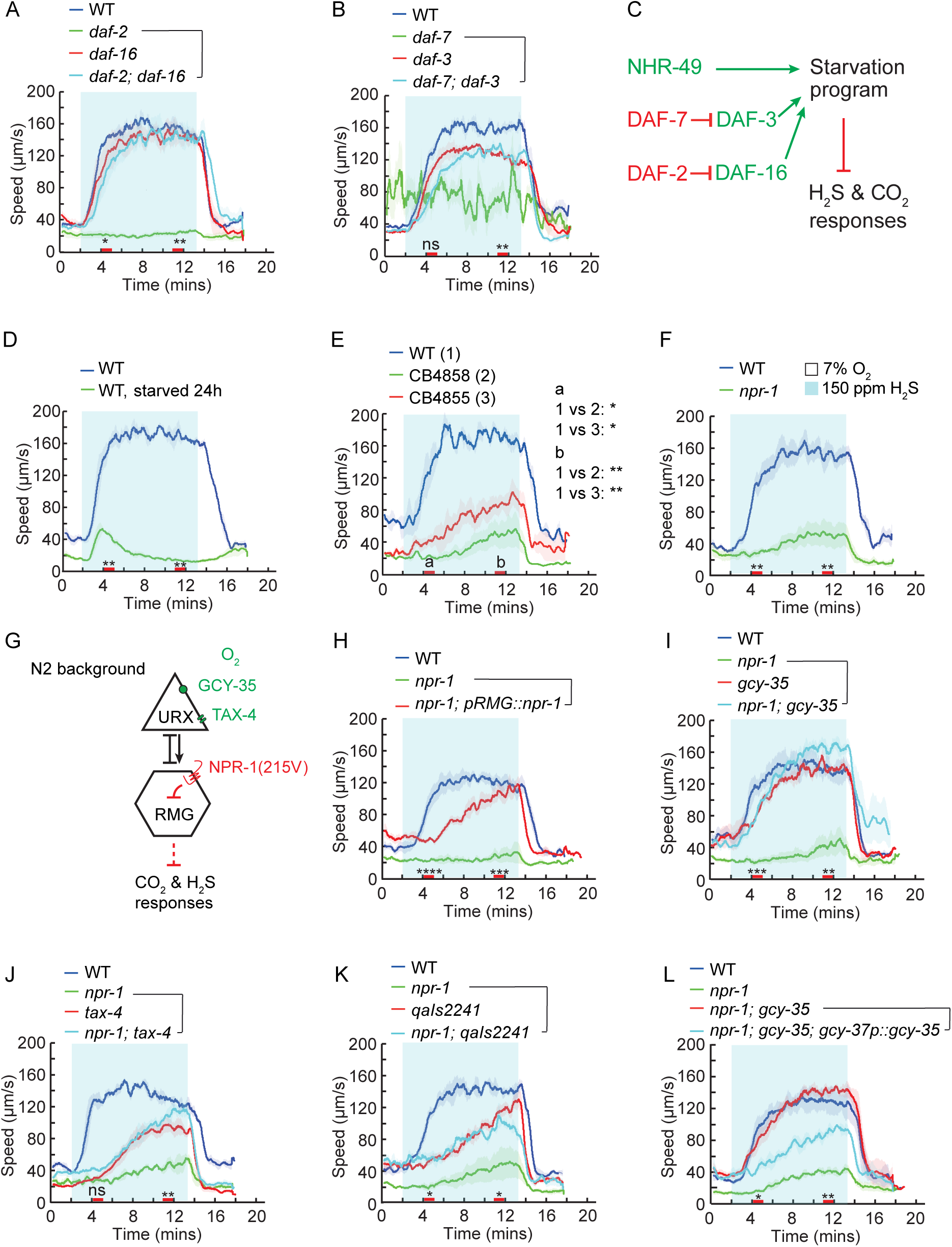
Insulin, TGF-β, and high O_2_ signaling pathways antagonizes H_2_S avoidance. (**A, B**) Locomotory speed responses to a switch from 7% O_2_ to 150 ppm H_2_S balanced with 7% O_2_ for animals of the indicated genotype: WT, *daf-2(e1370), daf-16(mgDf47)*, and *daf-2(e1370); daf-16(mgDf47)* double mutants (**A**); WT, *daf-7(e1372), daf-3(mgDf90),* and *daf-7(e1372); daf-3(mgDf90)* double mutants (**B**). For the *daf-2* assays, WT and *daf-2* mutants were maintained at 15°C. L4 animals were picked and shifted to 25°C until day-one adults, then assayed at room temperature. (**C**) Hypothetical model for the regulation of H_2_S avoidance by a starvation program involving the insulin, TGF-β, and NHR-49 pathways. (**D**) Locomotory speed response to a switch from 7% O_2_ to 150 ppm H_2_S balanced with 7% O_2_ in fed and starved WT animals. (**E, F**) Locomotory speed responses to a switch from 7% O_2_ to 150 ppm H_2_S balanced with 7% O_2_ for animals of the indicated genotype: WT and wild isolates (CB4858 and CB4855) (**E**); WT and *npr-1(ad609)* (**F**). (**G**) Hypothetical model showing how O_2_ regulates H_2_S avoidance via the inhibitory RMG circuit activity, which is modulated by O_2_ sensory inputs and NPR-1 signaling. (**H–L**) Locomotory speed responses to a switch from 7% O_2_ to 150 ppm H_2_S balanced with 7% O_2_ for animals of the indicated genotype: WT, *npr-1(ad609)*, and transgenic *npr-1(ad609)* expressing *npr-1* in RMG neurons using Cre-LoxP system (Macosko et al., 2009) (**H**); WT, *npr-1(ad609), gcy-35(ok769)*, and *npr-1(ad609); gcy-35(ok769)* double mutants (**I**); WT, *npr-1(ad609), tax-4(p678)* and *npr-1(ad609); tax-4(p678)* double mutants (**J**); WT, *npr-1(ad609), qaIs2241*(genetic ablation of AQR, PQR and URX neurons) and *npr-1(ad609); qaIs 2241* (**K**); WT, *npr-1(ad609), npr-1(ad609); gcy-35(ok769)* double mutants and *transgenic npr-1(ad609); gcy-35(ok769)* double mutants expressing *gcy-35* cDNA under *gcy-37* promoter, which drives *gcy-35* expression in O_2_ sensing neurons (**L**). For the comparison of acute locomotory responses between strains, red bars on the x-axis represent two intervals (4– 5 minutes and 11–12 minutes, labeled a and b, respectively, in panel E) used for statistical analysis. **** = *p* < 0.0001, *** = *p* < 0.001, ** = *p* < 0.01, * = *p* < 0.05, ns = not significant, Mann–Whitney U test.

### The O_2_ sensing circuit antagonizes H_2_S avoidance in wild isolates

Wild strains of *C. elegans* thrive in environmental niches characterized by variable O_2_, CO_2_ and H_2_S levels (Adams et al., 1979; Bretscher et al., 2008; Budde & Roth, 2011; Gea, Barrena, Artola, & Sanchez, 2004; Hallem & Sternberg, 2008; Morra & Dick, 1991; Oshins, Michel, Louis, Richard, & Rynk, 2022; Patange et al., 2025; Rodriguez-Kabana et al., 1965). Compared to the laboratory reference strain N2, wild isolates display robust responses to high O_2_ but reduced sensitivity to CO_2_ stimulation (Beets et al., 2020; Bretscher et al., 2008; Carrillo, Guillermin, Rengarajan, Okubo, & Hallem, 2013; Hallem & Sternberg, 2008; Kodama-Namba et al., 2013; McGrath et al., 2009). It is thought that the unique O_2_ and CO_2_ responses of N2 strain evolved during its domestication on solid media at atmospheric gas concentrations (Sterken, Snoek, Kammenga, & Andersen, 2015). We hypothesized that wild *C. elegans* strains might also exhibit an attenuated response to H_2_S, and therefore included a set of wild strains in our candidate gene screen. Consistent with this hypothesis, wild isolates including CB4855, CB4856, and CB4858 exhibited reduced locomotory speed in H_2_S (***Figure 3E, Figure 3— figure supplement 1J***).

A naturally occurring variation in the neuropeptide receptor NPR-1 predominantly determines the differences in locomotory responses to O_2_ and CO_2_ between the laboratory N2 strain and wild isolates. The NPR-1(215F) variant found in wild isolates is less active than the NPR-1(215V) found in N2 animals (de Bono & Bargmann, 1998). We wondered whether the variations in the *npr-1* gene also contribute to differences in the acute response to H_2_S. The absence of *npr-1* was sufficient to reduce the speed response to H_2_S, suggesting that active NPR-1 signaling promotes H_2_S avoidance (***Figure 3F***). NPR-1(215V) primarily acts within the RMG interneurons to modulate responses to environmental stimuli by inhibiting signal output from these neurons (***Figure 3G***) (Laurent et al., 2015; Macosko et al., 2009). Cell-specific expression of the highly active *npr-1(215V)* variant in RMG neurons effectively restored the H_2_S-evoked locomotory speed in *npr-1* null mutant worms (***Figure 3H***). Furthermore, optogenetic stimulation of RMG interneurons using channelrhodopsin-2 (ChR2) significantly dampened the locomotory response to H_2_S in N2 animals, suggesting that high RMG activity inhibits H_2_S responses (***Figure 3—figure supplement 1K***).

The perception of 21% O_2_ involves the soluble guanylate cyclases GCY-35/GCY-36 and the downstream cyclic nucleotide-gated (CNG) channels TAX-4/TAX-2 in the URX, AQR, and PQR O_2_-sensing neurons, which relay the signals to RMG (Busch et al., 2012; Cheung, Cohen, Rogers, Albayram, & de Bono, 2005; Couto, Oda, Nikolaev, Soltesz, & de Bono, 2013; Gray et al., 2004; Laurent et al., 2015; Persson et al., 2009; Zimmer et al., 2009). We investigated whether disrupting the O_2_ sensing machinery would have the same effect as inhibiting RMG signaling pathway by NPR-1(215V). Mutating *gcy-35* and *tax-4*, or genetically ablating the O_2_ sensing neurons URX, AQR, and PQR by cell-specific expression of the pro-apoptotic gene *egl-1* restored H_2_S-evoked speed response in *npr-1* null mutants (***Figure 3I–K***). Specific expression of *gcy-35* cDNA in O_2_ sensing neurons was sufficient to repress this response to H_2_S in *npr-1; gcy-35* double mutants (***Figure 3L***). Finally, enhancing the presynaptic activity of URX, AQR, PQR neurons by the expression of a gain-of-function PKC-1(A160E) (Dekker, Mcintyre, & Parker, 1993; Hiroki et al., 2022) significantly dampened H_2_S-induced locomotory speed (***Figure 3—figure supplement 1L***). Taken together, these observations indicate that activation of the O_2_-sensing circuit inhibits the neural pathway required for H_2_S-evoked avoidance response.

### H_2_S exposure reprograms gene expression in *C. elegans*

Wild isolates that were relatively insensitive to H_2_S exposure (***Figure 3E, Figure 3—figure supplement 1J***) likely evolved adaptations to persist in environments where H_2_S levels may transiently increase (Budde & Roth, 2011; Patange et al., 2025; Rodriguez-Kabana et al., 1965; Romanelli-Cedrez et al., 2024). These adaptations to H_2_S are mediated, at least in part, by the reprogramming of gene expression (Miller et al., 2011). To identify the genes whose expression was induced by prolonged H_2_S exposure, we conducted a comparative analysis of transcriptome profiles in animals exposed to 50 ppm or 150 ppm H_2_S for 1 hour, 2 hours, or 12 hours. RNA-seq analysis revealed that exposure to either 50 ppm or 150 ppm H_2_S for one hour was sufficient to trigger significant changes in gene expression, with 518 or 304 genes showing differential expression, respectively (***Figure 4A, Supplementary file 2***). Genes induced by H_2_S exposure for 1 or 2 hours displayed considerable overlap (***Figure 4B, Figure 4—figure supplement 1A***). Gene Ontology (GO) analysis revealed that biological processes such as defense against bacteria and cysteine biosynthesis from L-serine were significantly enriched after H_2_S exposure (***Figure 4C, D, Figure 4—figure supplement 1B, C***). As expected, we observed a robust induction of genes involved in H_2_S detoxification (***Figure 4E, F***) (Horsman & Miller, 2016; Miller et al., 2011; Niu et al., 2011; Vora et al., 2022). H_2_S is detoxified in mitochondria by sulfide:quinone oxidoreductase SQRD-1 to produce persulfide, which is then metabolized by sulfur dioxygenase ETHE-1 to generate sulfite. Thiosulfate transferase further oxidizes sulfite to produce thiosulfate (Hildebrandt & Grieshaber, 2008). Glutathione S-transferases (GSTs) remove accumulated sulfur molecules during H_2_S oxidation (Jackson, Melideo, & Jorns, 2012). Notably, *gst-19* and *sqrd-1* were among the most significantly upregulated genes after one or two hours of exposure to either 50 ppm or 150 ppm H_2_S, while *ethe-1* showed a weak increase (***Figure 4E, F, Figure 4—figure supplement 1D, E, Supplementary file 2***).

**Figure 4.**
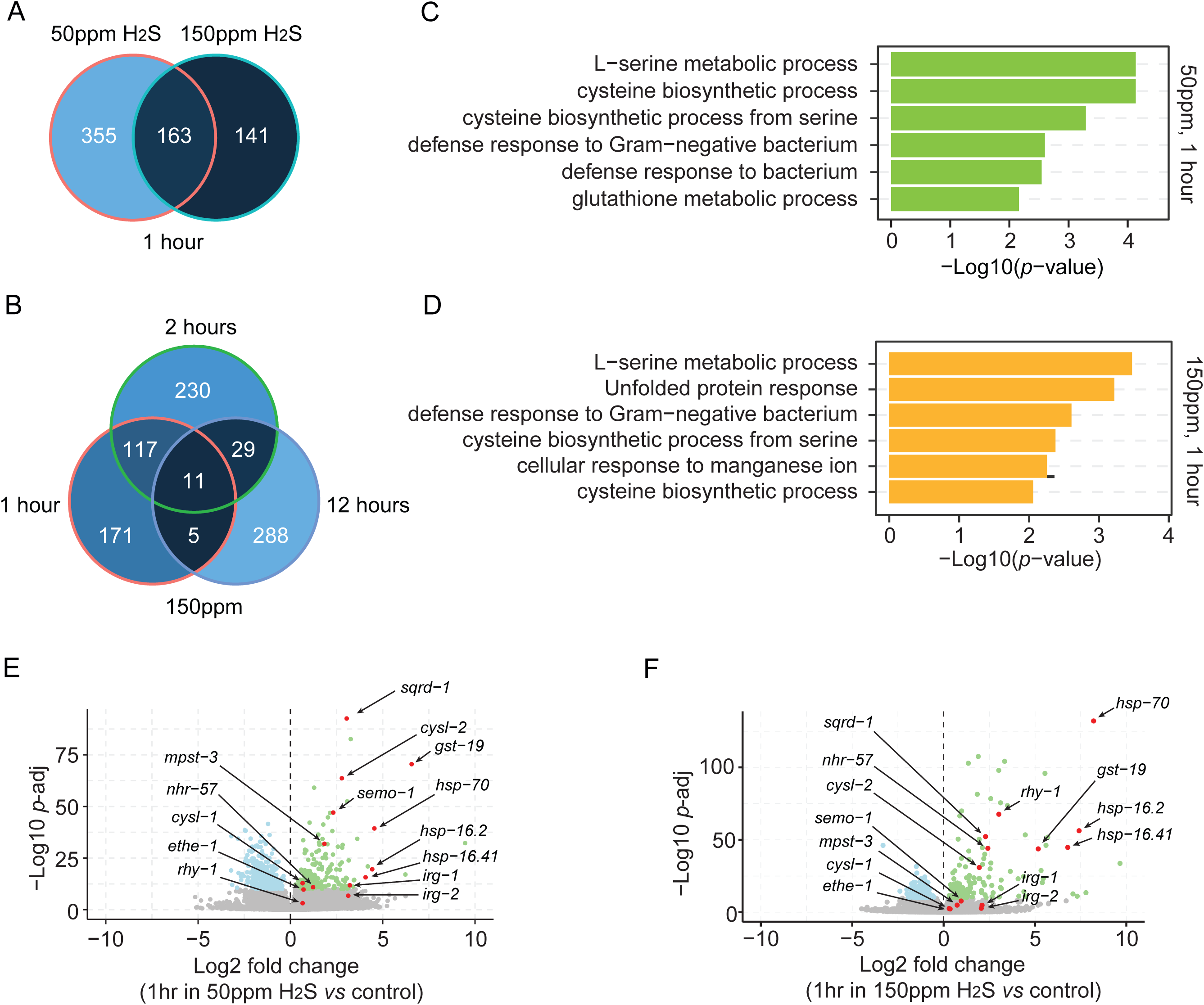
Prolonged H_2_S exposure reprograms gene expression in *C. elegans*. **(A)** Venn diagram displaying the number of differentially expressed genes after 1-hour exposure to 50 ppm or 150 ppm H_2_S balanced with 7% O_2_ in WT animals (adjusted *p* < 1e-10). **(B)** Venn diagram displaying the number of differentially expressed genes after 1-, 2-, and 12-hour exposure to 150 ppm H_2_S balanced with 7% O_2_ in WT animals (adjusted *p* < 1e-10). (**C, D**) Significantly enriched GO categories for differentially expressed genes with adjusted *p* < 1e-10 in WT animals exposed to 50 ppm (**C**) or 150 ppm (**D**) H_2_S balanced with 7% O_2_ for 1 hour. (**E, F**) Volcano plots showing the differentially expressed genes with adjusted *p* < 1e-10 in WT animals exposed to 50 ppm (**E**) or 150 ppm (**F**) H_2_S balanced with 7% O_2_ for 1 hour. A set of genes involved in H_2_S detoxification, cysteine metabolism and stress response were highlighted in red.

H_2_S exposure is known to activate the HIF-1 pathway (Budde & Roth, 2010, 2011; Horsman et al., 2019; Ma et al., 2012; Miller et al., 2011; Powell-Coffman, 2010; Topalidou & Miller, 2017). In line with this, a proportion of genes with HIF-1 regulated promoters displayed increased expression after 1 or 2 hours of H_2_S exposure (***Figure 4—figure supplement 1F***). Among these were the aforementioned detoxifying enzymes GST-19, SQRD-1, and ETHE-1, as well as CYSL-1, CYSL-2, and CDO-1, which are involved in cysteine metabolism (***Supplementary file 2***). Consistent with earlier studies, we also observed differential expression of a set of SKN-1 targets in response to chronic H_2_S exposure (***Figure 4—figure supplement 1G***) (Horsman et al., 2019; Miller et al., 2011; Niu et al., 2011). This included the heat shock protein genes *hsp-16.2* and *hsp-16.41* (***Figure 4E, F, Figure 4—figure supplement 1D, E***). Surprisingly, we detected a rapid increase in the expression of genes associated with intracellular H_2_S production, including the methanethiol oxidase-encoding gene *semo-1* and the mercaptopyruvate sulfurtransferase-encoding gene *mpst-3* (***Figure 4E, F, Figure 4—figure supplement 1D, E***) (Philipp et al., 2022; Qabazard et al., 2014). Elevated expression of 3-mercaptopyruvate sulfurtransferase has similarly been reported in sulfide spring fish, suggesting a conserved response mechanism to H_2_S (Kelley et al., 2016; Mathew, Schlipalius, & Ebert, 2011; Qabazard et al., 2014). However, the functional implications of increased expression of H_2_S synthesis genes remains unclear.

When exposed to H_2_S for 12 hours, the differentially expressed genes showed reduced overlap with those identified at earlier time points (***Figure 4B, Figure 4—figure supplement 1A***), suggesting that distinct defense strategies may be employed at various stages of H_2_S exposure. Furthermore, substantial overlap was observed between the sets of genes induced by 50 ppm and 150 ppm H_2_S after 1h or 2h exposure. However, this overlap decreased significantly after 12 hours exposure (***Figure 4—figure supplement 1H–J***). Additionally, we noticed that 150 ppm H_2_S specifically triggered the expression of the HIF-1 regulator *rhy-1* and the HIF-1 target *nhr-57* at all exposure time points, a response that was less pronounced at 50 ppm H_2_S (***Figures 4E, F, Figure 4—figure supplement 1D, E, K and L***). Together, these findings suggest that multiple defense mechanisms are induced to adapt to the stress imposed by H_2_S, which can diverge over time and at different H_2_S concentrations.

### Locomotory speed response to H_2_S is modulated by HIF-1-induced detoxification

Given that HIF-1 signaling is robustly activated shortly after H_2_S exposure (***Figure 4—figure supplement 1F***) (Budde & Roth, 2010, 2011; Miller et al., 2011), we next sought to determine how HIF-1 signaling might contribute to H_2_S-evoked avoidance response, and began by examining the effects of HIF-1 stabilization. Acclimating animals in 1% O_2_ for 12 hours or longer significantly decreased the speed response to H_2_S, indicating that HIF-1 stabilization inhibits the H_2_S-evoked avoidance behavior **(*Figure 5A*)**. To confirm this observation, we examined several mutants with stabilized HIF-1. The conserved proline-4-hydroxylase PHD/EGL-9 and the von Hippel-Lindau (VHL) tumor suppressor protein are required for HIF-1 degradation (Kaelin & Ratcliffe, 2008; Semenza, 2010). Disruption of either *egl-9* or *vhl-1* leads to HIF-1 stabilization under normoxic conditions. Similar to the effect of prolonged hypoxia, both mutants exhibited a markedly reduced locomotory speed in H_2_S **(*Figure 5B*).** Interestingly, the omega-turn and reversal responses to H_2_S were also substantially inhibited in *egl-9* mutants **(*Figure 5—figure supplement 1A, B*)**. In contrast, the locomotory speed response to 1% O_2_ remained largely unaffected in *egl-9* deficient animals **(*Figure 5—figure supplement 1C*),** suggesting a selective impairment of the H_2_S response. In addition, expressing the non-degradable forms of HIF-1, *hif-1*(P621A or P621G), under the pan-neuronal *unc-14* promoter or under the endogenous *hif-1* promoter, was sufficient to inhibit H_2_S evoked speed responses (***Figure 5C***). Together, these findings indicate that the activation of HIF-1 signaling during prolonged H_2_S or hypoxia exposure mediates an adaptive response associated with reduced behavioral responses to H_2_S, and that HIF-1 stabilization in neurons alone is sufficient to promote this behavioral adaptation.

**Figure 5.**
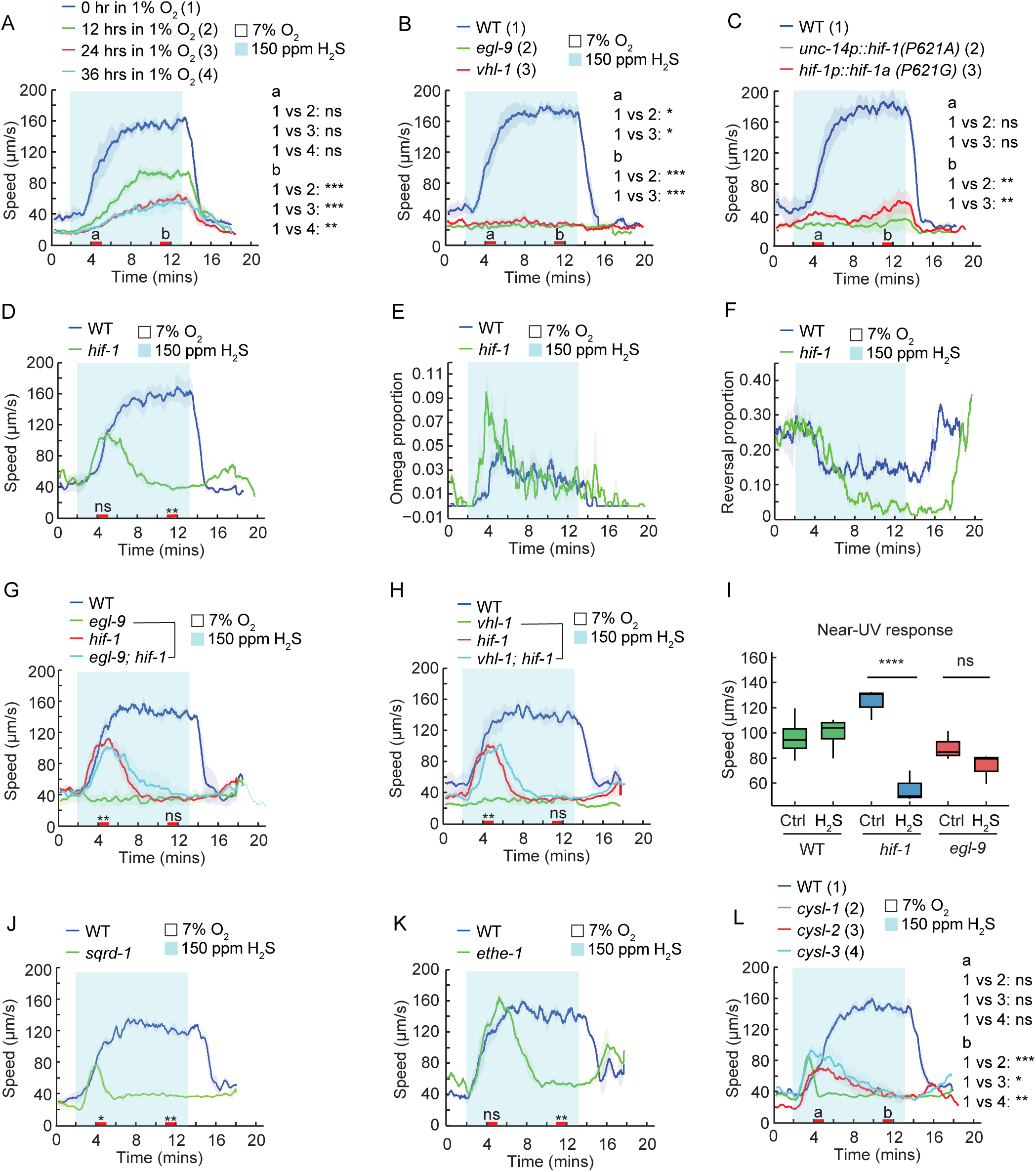
Acute response to H_2_S is modulated by HIF-1 signaling. (**A**) Locomotory speed responses of WT animals to a switch from 7% O_2_ to 150 ppm H_2_S balanced with 7% O_2_ after incubation of the animals for 0, 12, 24, or 36 hours in 1% O_2_. (**B, C**) Locomotory speed responses to a switch from 7% O_2_ to 150 ppm H_2_S balanced with 7% O_2_ for animals of the indicated genotype: WT, *egl-9(sa307)*, and *vhl-1(ok161)* (**B**); WT and transgenic animals expressing the non-degradable form of HIF-1 (P621A or P621G) under the pan-neuronal *unc-14* promoter or the endogenous *hif-1* promoter, respectively (**C**). (**D–F**) Avoidance responses of WT and *hif-1(ia4)* to a switch from 7% O_2_ to 150 ppm H_2_S balanced with 7% O_2_, including locomotory speed (**D**), reorientation (omega-turn) (**E**), and reversal (**F**). (**G, H**) Locomotory speed responses to a switch from 7% O_2_ to 150 ppm H_2_S balanced with 7% O_2_ for animals of the indicated genotype: WT, *egl-9(sa307), hif-1(ia4)*, and *egl-9(sa307); hif-1(ia4)* double mutants (**G**); WT, *vhl-1(ok161), hif-1(ia4)*, and *vhl-1(ok161); hif-1(ia4)* double mutants (**H**). (**I**) Average locomotory speed during 2 minutes of exposure to near-UV light for WT, *hif-1(ia4)*, and *egl-9(sa307)* before (Ctrl = control) and after 30 minutes of exposure to 150 ppm H_2_S balanced with 7% O_2_. (**J–L**) Locomotory speed responses to a switch from 7% O_2_ to 150 ppm H_2_S balanced with 7% O_2_ for animals of the indicated genotype: WT and *sqrd-1(tm3378)* (**J**); WT and *ethe-1(yum2895)* (**K**); WT, *cysl-1(ok762), cysl-2(ok3516)*, and *cysl-3(yum4)* (**L**). For the comparison of acute locomotory responses between strains, red bars on the x-axis represent two intervals (4–5 minutes and 11–12 minutes, labeled a and b, respectively, in panels A, B, C, and L) used for statistical analysis. *** = *p* < 0.001, ** = *p* < 0.01, * = *p* < 0.05, ns = not significant, Mann–Whitney U test.

On the other hand, animals lacking HIF-1 signaling have previously been shown to display increased sensitivity to H_2_S and become paralyzed rapidly upon exposure (Budde & Roth, 2010; Horsman et al., 2019). Consistent with this, *hif-1* mutants exhibited a brief increase in speed upon H_2_S exposure, which rapidly declined to baseline levels (***Figure 5D***). They also displayed a stronger initial omega-turn response and greater inhibition of the reversal rate than WT (***Figure 5E, F***), confirming that animals deficient in HIF-1 signaling are sensitized to H_2_S exposure. This transient avoidance behavior was also observed in *egl-9; hif-1* and *vhl-1; hif-1* double mutants (***Figure 5G, H***), suggesting that this phenotype is linked to the absence of HIF-1 signaling. Importantly, *hif-1* mutants displayed a robust speed response to other stimuli, such as near-UV light and 1% O_2_ stimulation ***(Figure 5I, Figure5—figure supplement 1D)***, suggesting that the brief avoidance response is specific to H_2_S rather than reflecting a general locomotory defect. Further supporting increased sensitivity to H_2_S in *hif-1* mutants, the locomotory speed response of *hif-1* mutants to subsequent near-UV light stimulation was nearly abolished after 30 minutes of H_2_S exposure, whereas *egl-9* mutants and WT animals remained fully responsive ***(Figure 5I)***. The reduced movement and lack of responsiveness to other stressors in *hif-1* mutants resemble the stress-induced sleep-like behavior previously observed in response to noxious heat (Byrne Rodgers & Ryu, 2020). Prompted by this observation, we next explored whether HIF-1-induced detoxification genes were also required to maintain H_2_S-evoked behavioral responses (***Figure 5—figure supplement 1E)***. Similar to *hif-1* mutants, *sqrd-1* and *ethe-1* mutants displayed brief responses to H_2_S, with locomotory activity rapidly returning to baseline levels ***(Figure 5J, K, Figure 5—figure supplement 1F–I)***. However, both mutants responded normally to 1% O_2_, suggesting a specific hypersensitivity to H_2_S toxicity ***(Figure 5—figure supplement 1J).*** Furthermore, we examined the contribution of additional genes whose expression was upregulated in response to H_2_S, including the sulfhydrylase/cysteine synthase-encoding genes *cysl-1*, *cysl-2*, and *cysl-3*, as well as *semo-1*. Disrupting any of these genes phenocopied the responses observed in *hif-1, ethe-1*, or *sqrd-1* mutants, while they displayed normal responses to 1% O_2_ **(*Figure 5L, Figure 5—figure supplement 1D, K–P***). These observations suggest that while the HIF-1-induced H_2_S detoxification system is not essential for initiating the response to H_2_S, it is required to maintain animals’ locomotory activity in the presence of H_2_S.

### Labile iron pool sustains the locomotory activity in H_2_S

Among the most differentially regulated genes upon exposure to H_2_S, we observed consistent downregulation of *ftn-1* and to a lesser extent *ftn-2*, whereas *smf-3* expression was increased under specific H_2_S conditions (***Figure 6A, B, Figure 6—figure supplement 1A–D***). The ferritin-encoding genes *ftn-1* and *ftn-2,* along with the iron transporter *smf-3*, are critical for maintaining intracellular labile iron homeostasis in *C. elegans* (Ackerman & Gems, 2012; Gourley, Parker, Jones, Zumbrennen, & Leibold, 2003; Rajan et al., 2019; Romney, Newman, Thacker, & Leibold, 2011), suggesting that H_2_S exposure may interfere with cellular iron storage. It has been shown that disrupting *ftn-1* increases labile iron pools in the cytoplasm, while its overexpression reduces these levels (Anderson & Leibold, 2014; Romney et al., 2011). We found that *ftn-1* mutants had enhanced speed response to H_2_S, whereas *ftn-1* overexpression attenuated H_2_S-induced speed increase without damaging acute response to 1% O_2_ (***Figure 6C, D, Figure 6—figure supplement 1E)***. In addition, *smf-3* mutants, which impair iron uptake, exhibited a reduced speed response to H_2_S (***Figure 6D***). These results support the idea that H_2_S exposure disrupts iron homeostasis.

**Figure 6.**
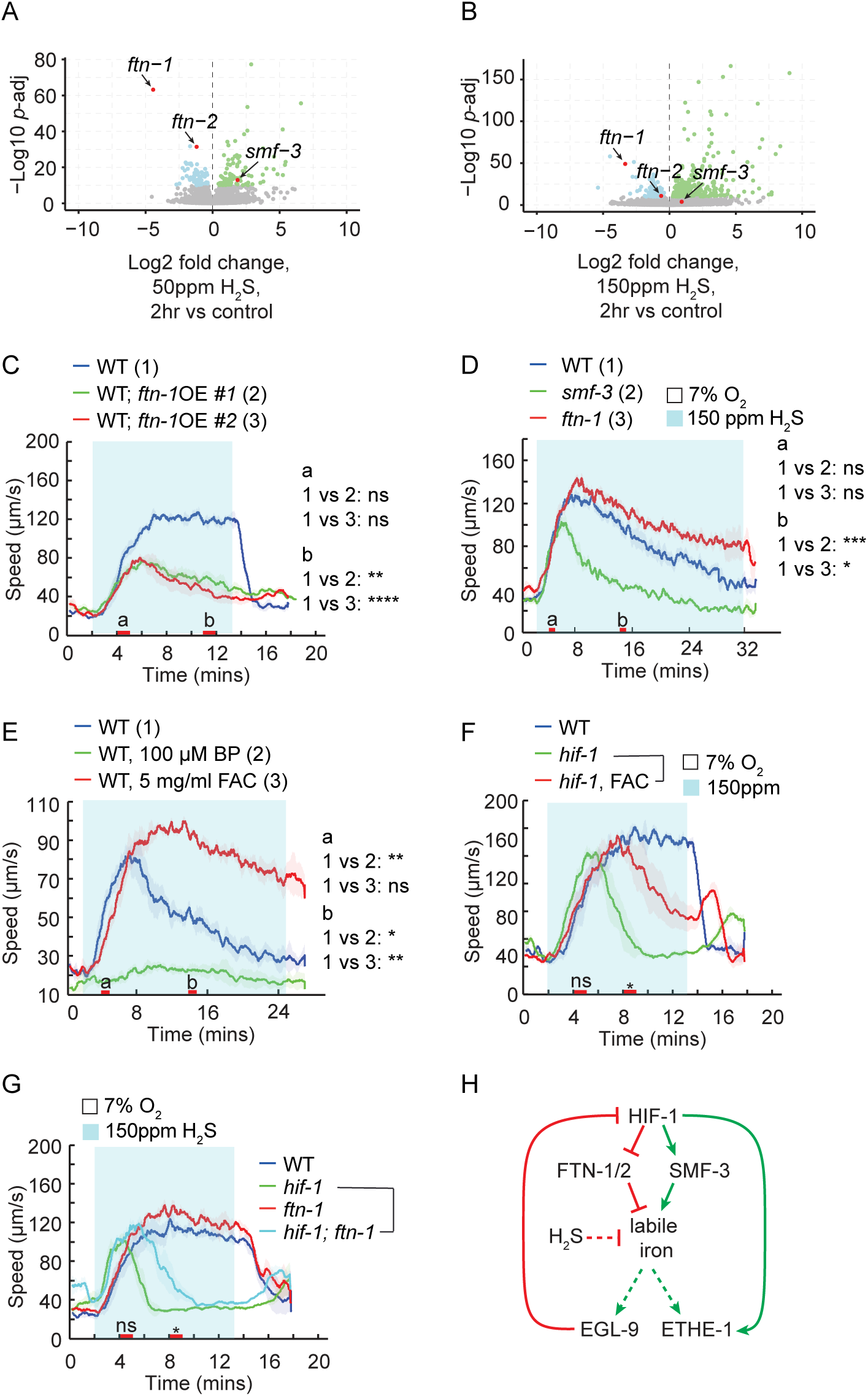
Labile iron pool sustains the locomotory activity in H_2_S. (**A, B**) Volcano plots showing the relative expression of the genes involved in the regulation of iron homeostasis in WT animals after 2-hour exposure in 50 ppm (**A**) or 150 ppm (**B**) H_2_S balanced with 7% O_2_. (**C–G**) Locomotory speed responses to a switch from 7% O_2_ to 150 ppm H_2_S balanced with 7% O_2_ for animals of indicated genotypes or treatments: WT and animals overexpressing *ftn-1* genomic DNA under its own promoter (#1 and #2 indicate two independent lines) (**C**); WT, *smf-3(ok1305)*, and *ftn-1(ok3625)* (**D**); WT, and WT pretreated with 100 μM 2,2′-Bipyridyl (BP) or with 5 mg/ml ferric ammonium citrate (FAC) in the presence of food for 16 hours (**E**); WT, *hif-1(ia4)*, and *hif-1(ia4)* mutants pretreated with 5 mg/ml ferric ammonium citrate (FAC) for 16 hours (**F**); WT, *hif-1(ia4), ftn-1(ok3625)*, and *hif-1(ia4); ftn-1(ok3625)* double mutants (**G**). (**H**) Hypothetical model of the interactions between labile iron pool and HIF-1 signaling. For the comparison of acute locomotory speed responses between strains or conditions, red bars on the x-axis represent two intervals (4–5 minutes and 11–12 minutes, labeled a and b, respectively, in panels C, D, and E) used for statistical analysis. **** = *p* < 0.0001, *** = *p* < 0.001, ** = *p* < 0.01, * = *p* < 0.05, ns = not significant, Mann– Whitney U test.

To complement our genetic analysis, we pharmacologically depleted labile irons. In the presence of the iron chelator 2,2’-dipyridyl (BP), the H_2_S-evoked speed response was fully lost (***Figure 6E***), whereas locomotory speed in response to hypoxia was maintained (***Figure 6—figure supplement 1F).*** We noticed that, similar to *hif-1* mutants, both *smf-3* mutants and BP-treated animals exhibited a strong initial omega-turn response to H_2_S (***Figure 6—figure supplement 1G, I)***, suggesting that these animals are hypersensitive to H_2_S toxicity. One possibility is that low iron levels compromise H_2_S detoxification capacity, as the detoxification enzyme ETHE-1 requires iron for its activity (Kabil & Banerjee, 2012; Pettinati, Brem, McDonough, & Schofield, 2015). By contrast, increasing iron levels by feeding ferric ammonium citrate (FAC) or by disrupting *ftn-1* significantly sustained animals’ locomotory speed in H_2_S (***Figure 6D, E***), likely by delaying H_2_S-mediated iron depletion. However, the reorientation and reversals were not significantly affected (***Figure 6—figure supplement 1G– J***). This impact of FAC supplementation and *ftn-1* disruption on speed response was largely lost in the absence of *hif-1* (***Figure 6F, G***), suggesting that the effects of iron require HIF-1-dependent detoxification (***Figure 6H)***. Yet, increased iron availability modestly improved the locomotory activity of *hif-1* mutants in H_2_S (***Figure 6F, G***), presumably by compensating for the low iron levels in *hif-1* mutant animals (Rajan et al., 2019; Romney et al., 2011).

### Mitochondrial function modulates H_2_S-evoked locomotion responses

The mitochondrial electron transport chain (ETC) is not only the primary target of H_2_S toxicity, but is also an essential component of the H_2_S detoxification pathways (Cooper & Brown, 2008; Khan et al., 1990; Nicholls & Kim, 1982; Romanelli-Cedrez et al., 2024). Based on this, we hypothesized that ETC plays a central role in the locomotory response to H_2_S. Indeed, mutating the genes critical for ETC function, including *gas-1*, *clk-1*, *mev-1,* and *isp-1*, impaired the H_2_S-evocked speed and reversal responses (***Figure 7A–D, Figure 7— figure supplement 1A–D)***. These data suggest either that dysfunctional ETC induces an adaptive response to cellular stressors or that a functional ETC is required to trigger and support H_2_S-evoked avoidance behavior. Consistent with the later model, exposing animals to rotenone, an ETC complex I inhibitor, robustly inhibited the speed responses to H_2_S at all time points we tested (***Figure 7E***). Similar to ETC mutants, WT animals exposed to rotenone for 2 hours also showed reduced basal locomotory activity (***Figure 7E***).

**Figure 7.**
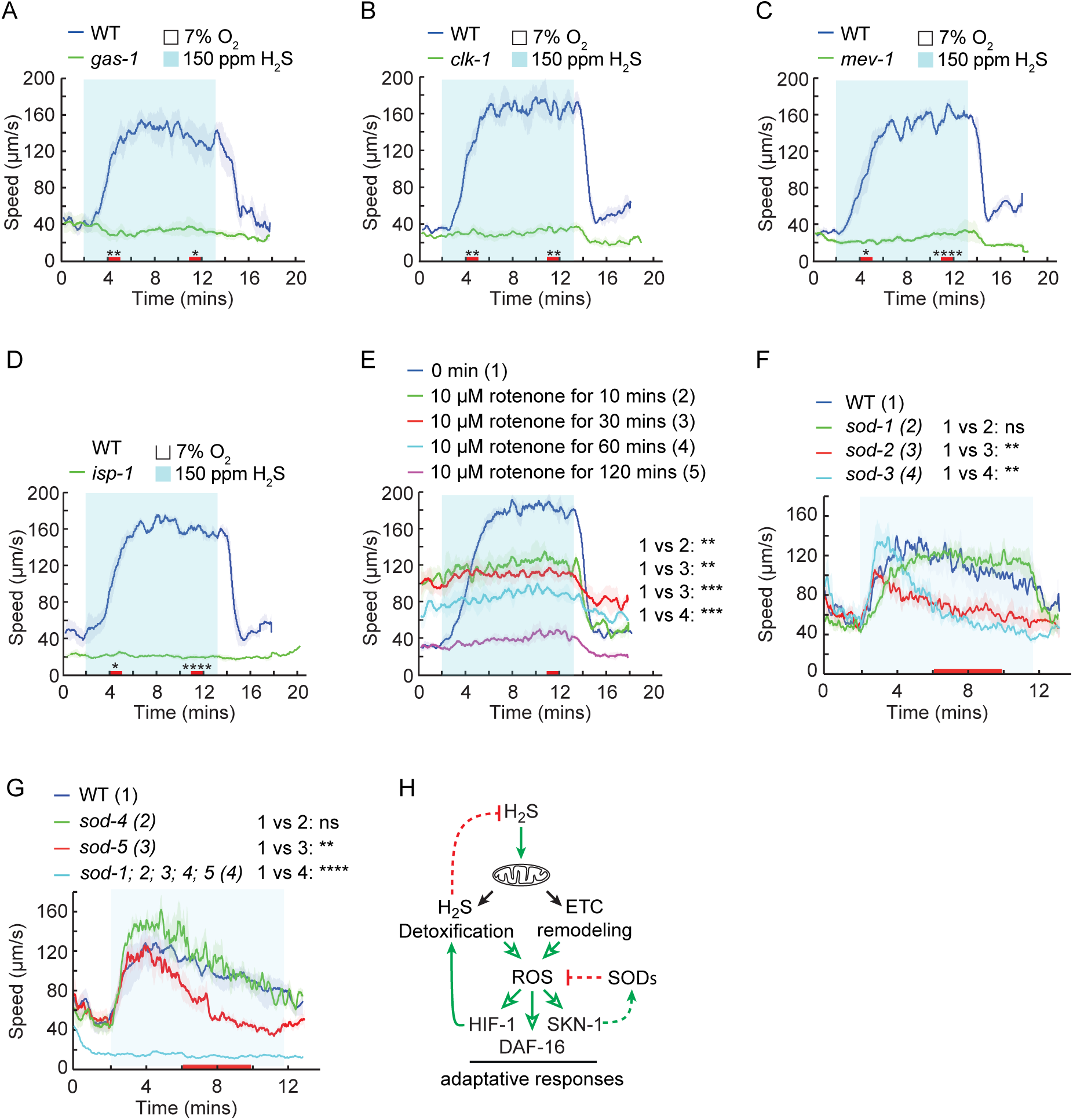
Mitochondrial function is required for acute response to H_2_S. (**A–D**) Locomotory speed responses to a switch from 7% O_2_ to 150 ppm H_2_S balanced with 7% O_2_ for animals of the indicated genotype: WT and *gas-1(fc21)* (**A**); WT and *clk-1(qm30)* (**B**); WT and *mev-1(kn1)* (**C**); WT and *isp-1(qm150)* (**D**). (**E**) Locomotory speed responses to a switch from 7% O_2_ to 150 ppm H_2_S balanced with 7% O_2_ for WT animals pretreated with 10 μM rotenone for 0 minute, 10 minutes, 30 minutes, 60 minutes or 120 minutes. For the comparison of acute locomotory speed responses between strains or conditions, red bars on the x-axis represent two intervals (4–5 minutes and 11–12 minutes) used for statistical analysis. *** = *p* < 0.001, ** = *p* < 0.01, * = *p* < 0.05, ns = not significant, Mann–Whitney U test. (**F– G**) Locomotory speed responses to a switch from 7% O_2_ to 150 ppm H_2_S balanced with 7% O_2_ for animals of the indicated genotype: WT, *sod-1(tm776)*, *sod-2(ok1030)*, and *sod-3(tm760)* (**F**); WT, *sod-4(gk101)*, *sod-5(tm1146)*, and *sod-1(tm783); sod-2(ok1030); sod-3(tm760); sod-4(gk101); sod-5 (tm1146)* (**G**). For the comparison of acute locomotory speed responses between strains or conditions, red bars on the x-axis represent the time interval (6–10 minutes) used for statistical analysis. *** = *p* < 0.001, ** = *p* < 0.01, * = *p* < 0.05, ns = not significant, Mann–Whitney U test. (**H**) Hypothetical model of the role of mitochondria in response to toxic levels of H_2_S. In this model, mitochondria play a dual role in H_2_S-evoked avoidance behavior. A transient burst of mitochondrial ROS triggered by high H_2_S levels initiates locomotory avoidance, whereas sustained ROS elevation activates stress-responsive pathways, including HIF-1, NRF2/SKN-1, and DAF-16/FOXO, promoting adaptation to prolonged H_2_S exposure.

In contrast to prolonged rotenone exposure, we noticed that transient rotenone exposure substantially increased animals’ basal locomotory speed (***Figure 7E***) (Onukwufor et al., 2022), suggesting that acute ETC interruption evokes behavioral avoidance, in a manner similar to that observed with other noxious stimuli. Since rotenone rapidly and persistently stimulates mitochondrial ROS production (Ochi et al., 2016; Ramsay & Singer, 1992; Zorov, Juhaszova, & Sollott, 2014), the transient locomotory increase during short rotenone exposure (***Figure 7E***) is likely driven by these ROS bursts, whereas reduced locomotory activity during prolonged exposure is presumably caused by chronically elevated ROS production. As toxic levels of H_2_S also induce ROS (Jia et al., 2020), this raised the possibility that H_2_S-evoked behavioral responses were modulated by the mitochondrial ROS. Therefore, we sought to further explore how mitochondrial ROS contributes to speed response to high H_2_S. We focused on superoxide dismutases (SODs), key enzymes in superoxide detoxification. The *C. elegans* genome encodes five *sod* genes: *sod-1* and *sod-5* encode cytosolic Cu/ZnSODs, *sod-2* and *sod-3* encode mitochondrial MnSODs, and *sod-4* encodes the extracellular Cu/ZnSOD isoforms (Doonan et al., 2008; Zubovych, Straud, & Roth, 2010). All *sod* single mutants exhibited a robust initial response to 150 ppm H_2_S ***(Figure 7F, G)*.** Interestingly, the mitochondrial SOD mutants *sod-2* and *sod-3*, as well as *sod-5*, showed an initial burst of locomotory activity followed by a rapid decline in speed in the presence of H_2_S ***(Figure 7F, G)***, but maintained a high locomotory speed in response to 1% O_2_ ***(Figure 7—figure supplement 1E)***. In the quintuple *sod-1; sod-2; sod-3; sod-4; sod-5* mutant, which lacks all superoxide dismutases and presumably accumulates higher ROS levels, the H_2_S-evoked speed and omega-turn responses were nearly abolished, and the proportion of animals exhibiting reversals remained consistently high, a pattern resembling that of *egl-9* and ETC mutants ***(Figure 7G, Figure 7—figure supplement 1F, G)***. Similar to *sod-2* and *sod-3*, the quintuple *sod* mutant exhibited robust speed response to 1% O_2_ (Zhao et al., 2022). In addition, the quintuple mutants responded normally to near-UV light after 30 minutes of H_2_S exposure (***Figure 7—figure supplement 1H)***, indicating that their neuromuscular function remains largely preserved. These observations suggest that either constitutively high ROS levels in quintuple mutants dampen H_2_S-evoked ROS transients, or they promote adaptation to H_2_S-induced stress, presumably via activation of stress-responsive pathways such as HIF-1, NRF2/SKN-1, and DAF-16/FOXO, ultimately reducing the locomotory responses (***Figure 7H***) (Lee, Hwang, & Kenyon, 2010; Lennicke & Cocheme, 2021; Patten, Germain, Kelly, & Slack, 2010).

## Discussion

Animals’ behavior and physiology are profoundly influenced by the environments in which they evolved. The laboratory strain N2, which is adapted to low atmospheric CO_2_ and H_2_S concentrations, robustly avoids both gases (Beets et al., 2020; Bretscher et al., 2008; Carrillo et al., 2013; Hallem & Sternberg, 2008; Kodama-Namba et al., 2013; McGrath et al., 2009). Upon acute exposure to H_2_S above 75 ppm, N2 exhibits an avoidance behavior characterized by reorientation and increased locomotion. This response is significantly reduced in wild isolates. These divergent responses between the laboratory strain and wild isolates are primarily driven by variations in the *npr-1* gene, which encodes a neuropeptide receptor, suggesting that *C. elegans* has undergone rapid evolutionary adaptation to its environment.

The behavioral response to a specific sensory stimulus is usually shaped by an interplay of multiple environmental and physiological cues. We observe that conditions dampening *C. elegans’* response to CO_2_ also impair the response to H_2_S. Specifically, the speed response to H_2_S is significantly inhibited at high O_2_ levels, and in mutants with defective insulin or TGF-β pathway that signal starvation. However, despite these shared modulations, CO_2_ and H_2_S responses involve distinct molecular mechanisms. For example, CO_2_ responses are mediated by the guanylate cyclase GCY-9 in BAG neurons (Hallem et al., 2011), which is dispensable for H_2_S responses. In addition, acute exposure to H_2_S induces a delayed acceleration compared to CO_2_. These data suggest that while responses to the two gases share a downstream neural circuit that can be dynamically modulated by the state of O_2_ sensing circuit, they are triggered by distinct mechanisms.

In a candidate gene survey, we excluded the potential involvement of globins, K^+^ channels, biogenic amines, and most guanylate cyclases in H_2_S responses. However, we observed that H_2_S avoidance requires the activity of guanylate cyclase DAF-11 in ASJ neurons, as well as neurosecretion from these neurons. Since the DAF-11 pathway and neurosecretion from ASJ neurons regulate developmental programs that modify sensory functions in *C. elegans* (Murakami, Koga, & Ohshima, 2001), it is not surprising that *daf-11* mutants display pleiotropic phenotypes including impaired H_2_S and CO_2_ responses (Hallem & Sternberg, 2008). In addition, we did not detect an H_2_S-evoked calcium transient in ASJ neurons. Therefore, although ASJ neurons and DAF-11 activity are clearly required, how H_2_S avoidance is triggered remains to be elucidated.

Consistent with previous reports (Horsman et al., 2019; Miller et al., 2011), H_2_S exposure substantially alters the gene expression in *C. elegans*, including those involved in H_2_S detoxification and iron homeostasis pathways. The repression of *ftn-1* and induction of *smf-3* observed during H_2_S exposure suggest that H_2_S depletes intracellular labile iron. Iron is a critical cofactor required for the activity of many enzymes and supports a wide range of cellular processes (Dev & Babitt, 2017; Hentze, Muckenthaler, & Andrews, 2004). One important enzyme that requires labile iron as cofactor is PHD/EGL-9, which targets HIF-1 for degradation. Changes in iron availability are therefore expected to modulate EGL-9 activity, which in turn alter the expression of HIF-1-regulated genes (Liochev, 1996; Myllyharju & Kivirikko, 1997; Read, Bentley, Archer, & Dunham-Snary, 2021; Xu & Moller, 2011). However, ETHE-1, a key H_2_S detoxification enzyme upregulated upon HIF-1 stabilization, also requires iron for its persulfide dioxygenase activity (Kabil & Banerjee, 2012; Pettinati et al., 2015). Therefore, H_2_S-evoked responses under varying iron availability are determined by the combined effects of labile iron on HIF-1 signaling and on the H_2_S detoxification enzymes, among other factors. For instance, although reduced iron availability in *smf-3* or BP-treated animals may promote the HIF-1 induced detoxification pathways by inactivating EGL-9, the actual enzymatic clearance of H_2_S is impaired due to low ETHE-1 activity. Supporting this, *smf-3* mutants and BP-treated animals display increased sensitivity to H_2_S, including enhanced initial omega-turn responses and rapid inhibition of locomotion. In contrast, increased iron availability in *ftn-1* mutants or FAC-treated animals likely delays iron depletion during H_2_S exposure, preserving ETHE-1 detoxification capacity and postponing the onset of HIF-1-mediated adaptations that are associated with reduced locomotion. This may explain the sustained high locomotory speed observed in *ftn-1* mutants or FAC-treated animals. In the absence of HIF-1, iron supplementation only partially improves the locomotory response to H_2_S, likely because the H_2_S detoxification system including ETHE-1 cannot be transcriptionally induced. The modest effect observed may instead reflect correction of iron deficiency in *hif-1* mutants.

At concentrations below 50 ppm, H_2_S is well tolerated or even preferred, with beneficial effects such as improved thermotolerance (Fawcett et al., 2015; Miller & Roth, 2007; Qabazard et al., 2014). Notably, behavioral avoidance is absent below 50 ppm H_2_S, suggesting that escape behavior is triggered only when H_2_S is not efficiently detoxified. This led us to hypothesize that animals with enhanced detoxification capacity would show reduced avoidance of otherwise toxic H_2_S levels. Indeed, the speed response to H_2_S is attenuated when the detoxification program is upregulated under conditions of constitutive activation of HIF-1, such as in *egl-9* or *vhl-1* mutants, after prolonged exposure to 1% O_2_, or when HIF-1 signaling is specifically activated in neurons. These observations support the idea that the initial avoidance response to H_2_S is triggered by neuronal detection of acute toxicity, while the subsequent decline in speed reflects a rapid activation of adaptive mechanisms, particularly through stabilization of HIF-1. The fact that wild type animals remain responsive to other stimuli after prolonged H_2_S exposure suggests that reduced locomotory speed in H_2_S reflects active desensitization rather than general paralysis. However, persistently high H_2_S exposure is likely to exhaust cellular defense systems, leading to toxicity and paralysis. By contrast, the rapid loss of locomotion observed in *hif-1* mutants and detoxification-defective mutants is mediated by a mechanism distinct from adaptation. Their loss of responsiveness to other stimuli after H_2_S exposure suggests that these animals likely become sensitized and rapidly intoxicated by H_2_S due to impaired detoxification. Therefore, H_2_S exposure promotes a behavioral program that includes an initial reorientation and acceleration responses followed by a progressive adaptation driven by cellular detoxification processes. A similar behavioral program has been observed in *C. elegans* during noxious heat exposure, which induces short-term heat avoidance followed by long-lasting cytoprotective adaptation and a gradual reduction in avoidance (Byrne Rodgers & Ryu, 2020).

The interaction of H_2_S with mitochondrial ETC is multifaceted, acting as an electron donor at low concentrations and becoming a potent toxicant at high levels (Szabo et al., 2014), in part by promoting superoxide generation through complex IV inhibition and reverse electron transport at complex I (Cooper & Brown, 2008; Jia et al., 2020; Khan et al., 1990; Nicholls & Kim, 1982; Romanelli-Cedrez et al., 2024). We propose that mitochondria play a dual role in H_2_S-evoked locomotory avoidance. On one hand, the mitochondrial ETC contributes to H_2_S detoxification and promotes adaptation. On the other hand, toxic levels of H_2_S remodel the ETC, leading to increased ROS production, which may serve as a trigger for the avoidance response. Supporting the idea that acute mitochondrial ROS generation initiates avoidance of high H_2_S levels, short-term rotenone exposure, known to promote mitochondrial ROS formation (Ochi et al., 2016; Ramsay & Singer, 1992; Zorov et al., 2014), substantially increases locomotory speed (Onukwufor et al., 2022). Meanwhile, the speed response to high H_2_S is fully suppressed by rotenone. This inhibition could result either from excessive mitochondrial ROS generated by rotenone, which may dampen the H_2_S-triggered ROS spike, or from direct complex I inhibition, which may disrupt other H_2_S signaling processes required to initiate avoidance. However, persistent mitochondrial ROS production appears to suppress high locomotory speed and inhibit responsiveness to H_2_S, as observed after 2-hour rotenone exposure, in mitochondrial ETC mutants, and in animals lacking all superoxide dismutases. One likely explanation is that mitochondrial ROS can activate a variety of stress-responsive pathways, including HIF-1, NRF2/SKN-1, and DAF-16/ FOXO signaling (Lee et al., 2010; Lennicke & Cocheme, 2021; Patten et al., 2010), priming animals’ adaptation to prolonged stress rather than causing toxicity. This is supported by the observation that even though SOD-deficient animals do not display strong initial locomotory responses to H_2_S, they remain responsive to other stimuli after 30 minutes of H_2_S exposure, suggesting that high ROS levels do not compromise general viability or the H_2_S detoxification capacity. Therefore, we favor a model in which mitochondrial ROS exert a biphasic effect on H_2_S-induced avoidance, facilitating H_2_S avoidance under acute conditions, and contributing to locomotory inhibition when it is chronically elevated. Overall, lack of a clear sensory machinery, the slow increase of locomotory speed in H_2_S (***Figure 1D***), the rotenone-evoked speed responses, and the strong modulation of H_2_S responses by mitochondrial ETC inhibition suggest that H_2_S may not be directly perceived by *C. elegans*. Instead, acute avoidance to H_2_S is likely initiated by ROS-induced toxicity.

In summary, this study unveils a novel behavior of *C. elegans* in their avoidance to H_2_S. The speed response to H_2_S is intricately shaped by environmental context, integrating inputs from other sensory cues such as food availability and O_2_ levels. Our findings suggest that H_2_S-induced locomotion arises from acute remodeling of mitochondrial ETC and associated ROS production, while highlighting the vital role of the HIF-1-induced detoxification pathways and iron homeostasis in protecting against H_2_S-induced mitochondrial toxicity. Further work is needed to validate this model and to elucidate the neural circuits mediating the behavioral response to ROS-induced toxicity.

## Materials and Methods

### Strains

*C*. *elegans* were maintained using standard protocols (Brenner, 1974). Strains used in this study are listed in ***Supplementary file 1*** and ***Supplementary file 3***.

### Preparation of H_2_S gas mixture

Different H_2_S concentrations were created as previously described (Miller & Roth, 2007). Briefly, the H_2_S-containing gas mixture were prepared by diluting 5000 ppm H_2_S in nitrogen (N_2_) with 7% O_2_ balanced with N_2_. The gas flow was tuned by Sierra Smart-Trak 100 mass flow controllers. H_2_S concentration in the mixture was measured using two H_2_S detectors (MSA ALTAIR 2X gas detector for H_2_S and Clip Single Gas Detector, SDG, CROWCON). The pre-defined gas mixtures of 7% O_2_, 1% O_2_ and 5% CO_2_ with N_2_ were purchased from Air Liquide Gas AB. 5000 ppm H_2_S stock in N_2_ was obtained from Linde Gas AB. Gas mixtures in all experiments were hydrated before use.

### Molecular Biology

The Multisite Gateway system (Thermo Fisher Scientific, United States) was used to generate expression vectors. Promoters, including *gcy-37* (2.7kb), *trx-1* (1 kb), *ftn-1*(2 kb), *daf-11*(3 kb), *odr-3*(2.7 kb), *gpa-11*(3 kb), *sra-6*(3 kb), *odr-1*(2.4 kb), *flp-21*(4.1kb), and *ocr-2*(2.4 kb) were amplified from N2 genomic DNA and cloned into pDONR P4P1 using BP clonase. g*cy-35*, *tax-4,* and *pkc-1* cDNAs were amplified using the first strand cDNA library as the template, while *daf-11* and *ftn-1* genomic sequences were amplified using genomic DNA as the template. The entry clones of these genes pDONR 221 were generated using BP reaction. To generate the gain of function mutation of *pkc-1* (A160E), Q5 Site-Directed Mutagenesis Kit (NEB) was used according to manufacturer instructions. All expression vectors were generated using LR reaction. The primer sequences were displayed in ***Supplementary file 4***.

To generate transgenic animals, the *daf-11* expression vectors were injected at the concentration of 20 ng/μl supplemented with 50 ng/μl of a coelomocyte co-injection marker (*unc-122p::GFP)* and 30 ng/μl of 1kb DNA ladder. For *tax-4* gene, the injection mixtures were prepared using 40 ng/μl of *tax-4* expression vectors, 50 ng/μl of a coelomocyte co-injection marker and 10 ng/μl of 1kb DNA ladder. The rest of expression constructs were injected at 50 ng/μl together with 50 ng/μl of a coelomocyte marker.

### CRISPR/Cas9 genome editing

Genes were disrupted using CRISPR/Cas9 mediated genome editing as described (Dokshin, Ghanta, Piscopo, & Mello, 2018; Ghanta & Mello, 2020). The strategy involved the utilization of homology-directed insertion of custom designed single strand DNA template (ssODN), which had two homology arms of 35bp flanking the targeted PAM site. Between two homology arms, a short sequence containing a unique restriction enzyme cutting site as well as in-frame and out-of-frame stop codons, was included. The insertion of ssODN template would delete 16 bases of coding sequence, ensuring the proper gene disruption. To prepare the injection cocktail, 0.5 μl of Cas9 protein (IDT) was mixed with 5 μl of 0.4μg/μl tracrRNA (IDT, United States) and 2.8 μl of 0.4 μg/μl crRNA (IDT). The mixture was incubated at 37°C for at least 10 minutes before 2.2 μl of 1μg/μl ssODN (or 500 ng dsDNA) and 2 μl of 0.6 μg/μl *rol-6* co-injection marker were added. Nuclease-free water was used to bring the final volume to 20 μl. The injection mixture was centrifuged for 2 minutes before use.

### Behavioral assays

H_2_S evoked locomotion activity was monitored as described previously (Laurent et al., 2015; Zhao et al., 2022). Briefly, OP50 bacteria were seeded on the assay plates 16 hours before use. The border of bacterial lawn was removed using a PDMS stamp. For each assay, 25 to 30 day-one adult animals were picked onto assay plates, allowed to settle down for 15 minutes, and subsequently sealed within microfluidic chambers. A syringe pump (PHD 2000, Harvard Apparatus) was employed to deliver gas mixtures into the microfluidic chamber at a constant flow of 3 ml/min. The rapid switch between different gas mixtures were controlled by Telfon valves coupled with ValveBank Perfusion Controller (AutoMate Scientific). The locomotory activity at different gas mixtures were monitored using a high-resolution camera (FLIR) mounted on a Zeiss Stemi 508 microscope. Videos were captured at a rate of 2 frames per second, starting with a 2-minute recording in 7% O_2_, followed by 10 or 11 minutes in H_2_S, and ending with 7% O_2_. For the long-term recording in H_2_S, video capture commenced with a 2-minute period in 7% O_2_, followed by a duration of 148 minutes in 150 ppm H_2_S, and concluded with a final 2-minute interval in 7% O_2_. For each strain or condition, 3 to 4 replicates were performed.

Near-UV light experiments were conducted to assess whether animals remained responsive following a 30-minute incubation in 150 ppm H_2_S balanced with 7% O_2_. For each experiment, 6-minute videos were recorded. The first 2 minutes captured the baseline locomotion of the animals under white light, followed by 2 minutes of exposure to near-UV light (435 nm, 0.7 mW/mm^2^), and concluded with 2 minutes of white light. 3 to 4 replicates were performed, with approximately 15 animals recorded per replicate.

The H_2_S gradient experiment was performed in a PDMS chamber connected to a pump delivering gas at 0.5 ml/min. To establish a gradient ranging from 150 ppm to 0 ppm H_2_S, we pumped 150 ppm H_2_S balanced with 7% O_2_ into one side and 7% O_2_ into the opposite side. After 25 minutes, the number of animals in each of the 5 sections was counted (***Figure 1G***). Four replicates were performed, each using 30 day-one adult animals.

To assess the effects of rotenone on H_2_S evoked locomotory response, day-one adult animals were subjected to 10 μM rotenone for 10min, 30min, 1 hour and 2 hours on the drug containing plates. Subsequently, the rotenone-treated animals were assayed on the standard assay plates without the drug. To explore the impact of ferric ammonium citrate (FAC) (F5879, Sigma-Aldrich) and 2,2′-Bipyridyl (BP) (D216305, Sigma-Aldrich) on behavioral response to H_2_S, L4 animals were exposed to 100 μM FAC or 5 mg/ml BP for 16 hours in the presence of bacterial food. The FAC or BP treated animals were assayed on FAC or BP containing plates, respectively.

Optogenetic experiments were performed as previously outlined (Zhao et al., 2022). Transgenic L4 animals were exposed to 100 μM all-trans retinal (ATR) (R2500, Sigma) in the dark for 16 hours. The ATR-fed animals were assayed under continuous illumination of 70 μW/mm^2^ blue light, which was switched on at the start of recording at 7% O_2_. Blue light was emitted from an ultra-high-power LED lamp (UHP-MIC-LED-460, Prizmatix), and the light intensity was determined using a PM50 Optical Power Meter (ThorLabs). To minimize the effect of transmitted light on the transgenic animals, a long-pass optical filter was used to eliminate the lights with short wavelengths during picking. All videos of behavioral analysis were analyzed using a home-made MATLAB program Zentracker (https://github.com/wormtracker/zentracker). For all behavior analyses, at least three assays were performed for each strain with more than 80 worms in total.

### Ca^2+^ Imaging

L4 animals expressing the GCaMP6s sensor in ASJ neurons were picked 24 hours before imaging. The agarose pads were prepared 1 hour before the experiment. Animals were incubated in 15 µl of 10 mM levamisole for 40 minutes prior to imaging. Between 15 and 20 animals were immobilized using 10 mM levamisole diluted in a concentrated OP50 bacterial suspension and mounted on a 2% agarose pad in M9 buffer on a glass slide. The animals were then covered with a microfluidic chamber. Imaging lasted for 16 minutes. O_2_ (7%) was pumped into the microfluidic chamber during the first and last 4 minutes. From minute 4 to 12, a mixture of 7% O_2_ and 150 ppm H_2_S was pumped.

Imaging was performed on a Nikon AZ100 microscope equipped with a TwinCam adapter (Cairn Research, UK), fitted with two DMK 33U monochrome cameras (The Imaging Source, Germany). A 2x AZ Plan Fluor objective with 1x zoom was used, with an exposure time of 300 ms. Excitation of GCaMP and RFP was provided by a CoolLED pE-300 (CoolLED, UK). The TwinCam adapter was mounted with emission filters for green fluorescent protein (510/20 nm) and red fluorescent protein (590/35 nm), along with a DC/T510LPXRXTUf2 dichroic mirror. Imaging data were analyzed using Neuron Analyzer, a custom-written Matlab program for processing image stacks (code available at https://github.com/neuronanalyser/neuronanalyser).

### RNA extraction and sequencing

To obtain RNA samples for RNA-seq, thirty young adult animals were picked to each fresh plate and allowed to lay eggs for 2 hours, after which the adult animals were removed and eggs were allowed to develop into day-one adults. Day-one animals were subsequently challenged with H_2_S for three different time periods (1 hour, 2 hours and 12 hours). For each time period, five plates of day-one animals were exposed to 50 ppm H_2_S, 150 ppm H_2_S or 7% O_2_ as a control group. Subsequently, the animals were immediately collected and rinsed three times with M9 buffer. After washing, the worm pellet was frozen in liquid nitrogen. Animals were homogenized using Bullet Blender (Next Advance) in the presence of Qiazol Lysis Reagent and 0.5mm Zirconia beads at 4°C. RNA was prepared using RNeasy Plus Universal Mini Kit (Qiagen). Six independent RNA samples were prepared for each condition. The 2100 Bioanalyzer instrument (Agilent) was used for the RNA quality control with a microfluidic chip specific for RNA (Agilent RNA 6000 Nano Kit). Then 1 μg of qualified RNA samples in RNase-free ddH2O was used for library preparation. Library was constructed and sequenced by Novogene.

### RNA-seq analysis

The RNA-seq data were aligned using STAR v2.7.9a (Dobin et al., 2013) and gene expression counts extracted by featureCounts v2.0.3 (Liao, Smyth, & Shi, 2014). Differential expression was calculated using DESeq2 (Love, Huber, & Anders, 2014). GO analysis was performed using the EnrichR R package (Kuleshov et al., 2016) on genes having *p*.adj < 1e-10. For brevity, several highly similar GO categories were manually omitted (full EnrichR output available at Github repository).

## Data availability

Raw sequencing data has been deposited at ArrayExpress # E-MTAB-13296. The R code is available at https://github.com/henriksson-lab/ce_h2s_tc.

## Acknowledgments

We thank the Caenorhabditis Genetics Center (funded by NIH Office of Research Infrastructure Programs P40 OD010440) and the National BioResources Project Japan for strains.

## Figure supplement legends

**Figure 2—figure supplement 1.**
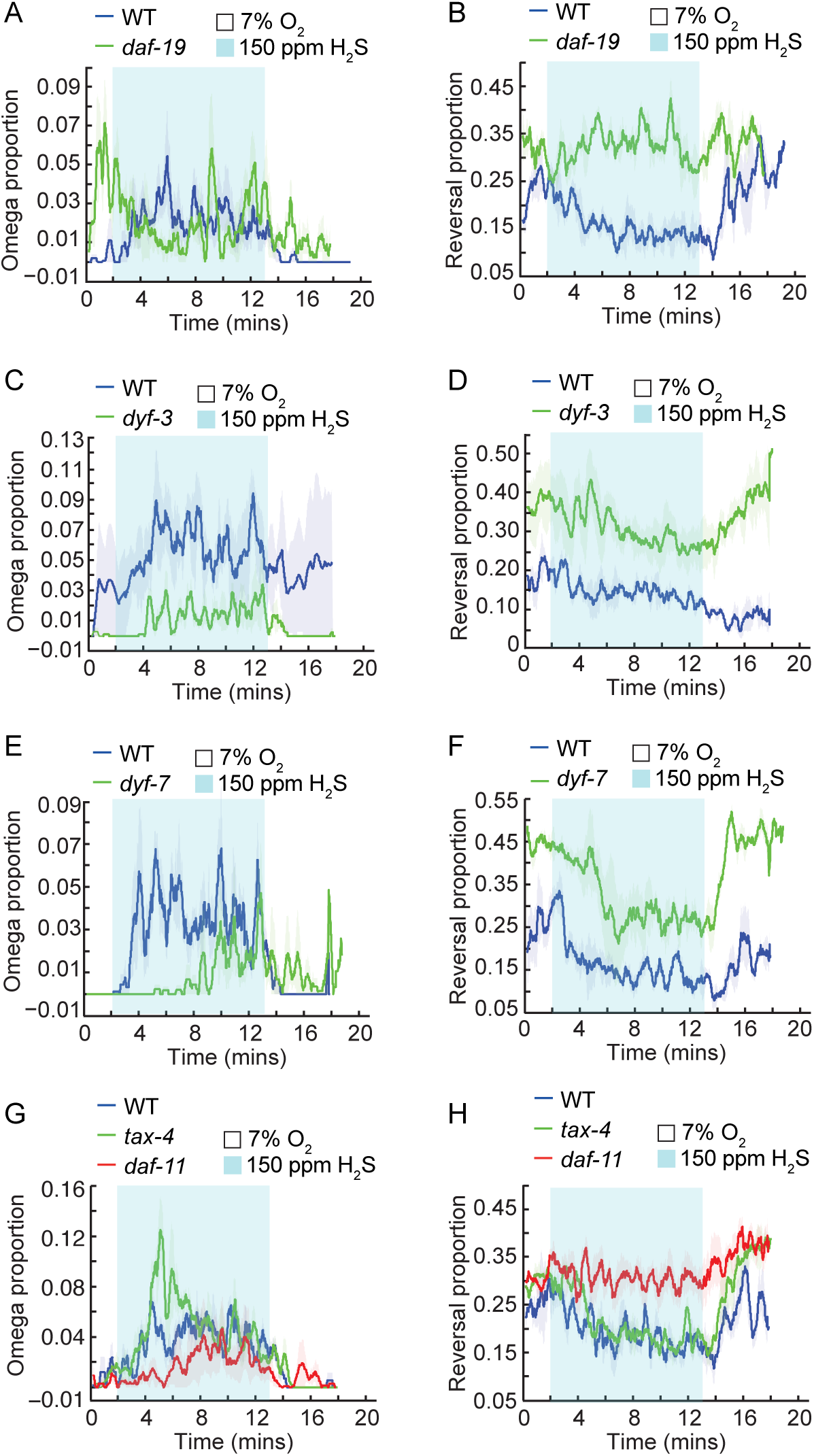
Acute avoidance response to H_2_S is regulated by cGMP signaling. (**A–H**) Fraction of animals undergoing reorientation movements (omega turns, left) or backward locomotion (reversals, right) during a switch from 7% O_2_ to 150 ppm H_2_S balanced with 7% O_2_ for animals of the indicated genotype: WT and *daf-19(m86)* (**A, B**); WT and *dyf-3(m185)* (**C, D**); WT and *dyf-7(m539)* (**E, F**); WT, *daf-11(m47)*, and *tax-4(p678)* (**G, H**).

**Figure 2—figure supplement 2.**
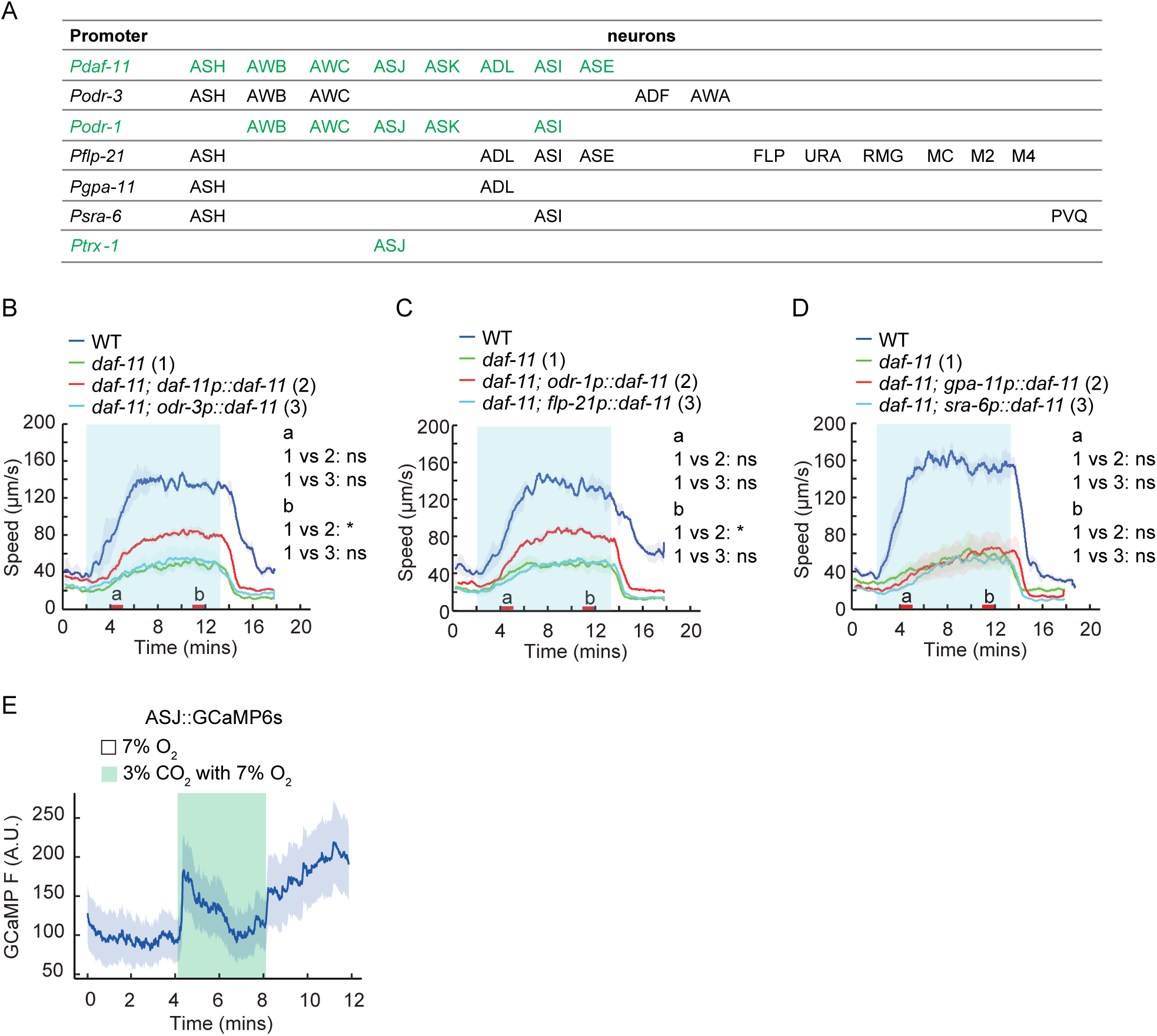
Acute response to H_2_S is regulated by *daf-11* signaling in ASJ neurons. (A) The detailed expression pattern of each promoter used to express *daf-11* genomic DNA. Promoters shown in green rescued the *daf-11(m47)* mutant phenotype. (**B–D**) Locomotory speed responses to a switch from 7% O_2_ to 150 ppm H_2_S balanced with 7% O_2_ for animals of the indicated genotype: WT, *daf-11(m47)*, and transgenic *daf-11* mutants expressing *daf-11* genomic DNA in different subsets of neurons. Red bars on the x-axis represent two intervals (4–5 minutes and 11–12 minutes, labeled a and b, respectively) used for statistical analysis. * = *p* < 0.05, ns = not significant, Mann–Whitney U test. (**E**) Calcium transients evoked in ASJ neurons of WT animals in response to a switch from 7% O_2_ to 3% CO_2_ balanced with 7% O_2_ for animals (N=16).

**Figure 3—figure supplement 1.**
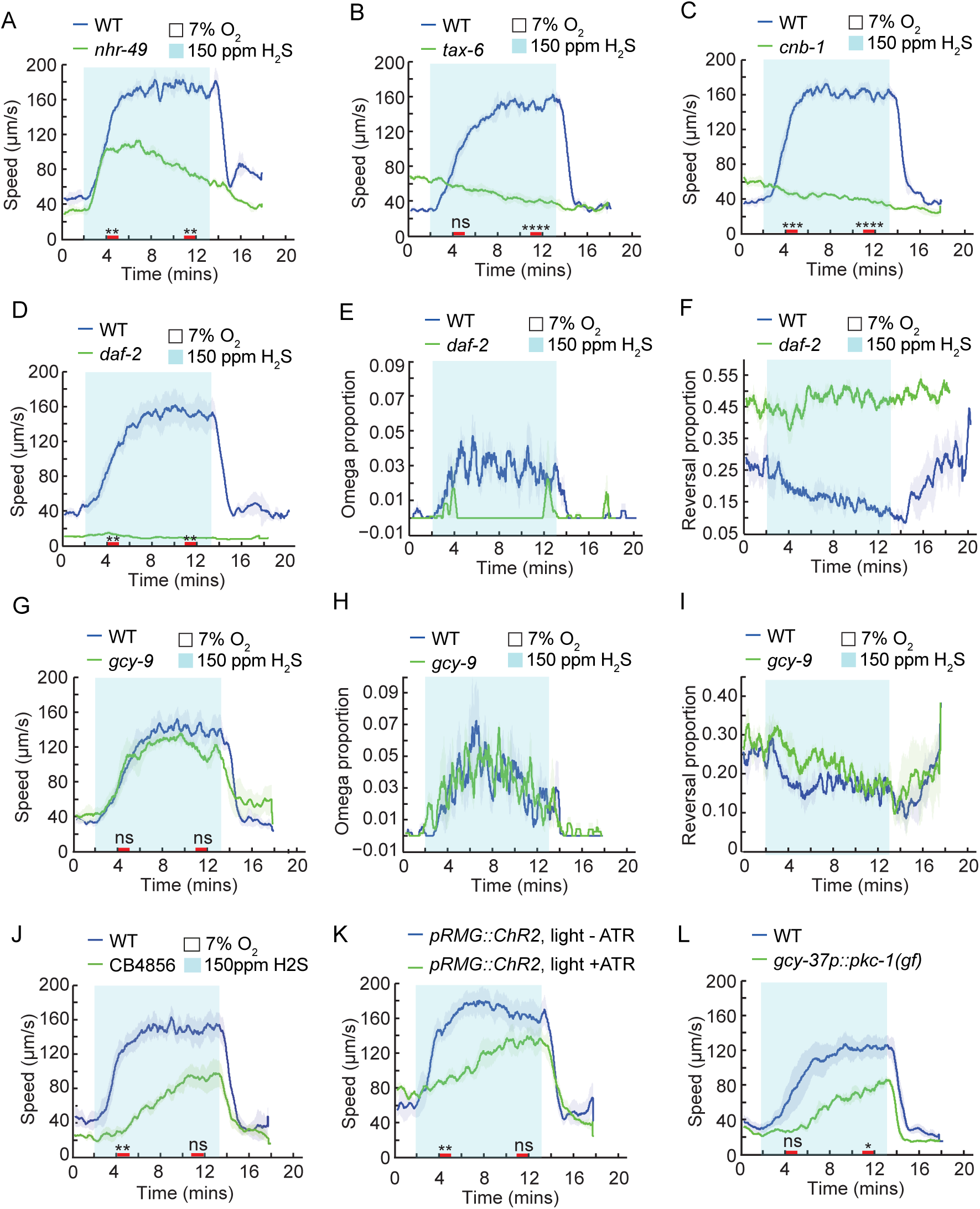
Insulin, TGF-β, and O_2_ signaling modulate locomotory response to H_2_S. (**A–C**) Locomotory speed responses to a switch from 7% O_2_ to 150 ppm H_2_S balanced with 7% O_2_ for animals of the indicated genotype: WT and *nhr-49(nr2041)* (**A**); WT and *tax-6(ok2065)* (**B**); WT and *cnb-1(ok276)* (**C**). (**D–F**) Locomotory responses to a switch from 7% O_2_ to 150 ppm H_2_S balanced with 7% O_2_ for WT and *daf-2(e1370)*, including locomotory speed (**D**), reorientation (omega turns) (**E**), and reversal (**F**). WT and *daf-2* mutants were maintained at 15°C. L4 animals were picked and shifted to 25°C until day-one adults, then assayed at room temperature. (**G–I**) Locomotory responses to a switch from 7% O_2_ to 150 ppm H_2_S balanced with 7% O_2_ for WT and *gcy-9(tm7632)*, including locomotory speed (**G**), reorientation (omega turns) (**H**), and reversal (**I**). (**J**) Locomotory speed responses to a switch from 7% O_2_ to 150 ppm H_2_S balanced with 7% O_2_ for WT animals and wild isolate CB4856. (**K**) Optogenetic stimulation of RMG neurons using channelrhodopsin-2 (ChR2). ChR2 was expressed in RMG neurons using the Cre-LoxP system (Macosko et al., 2009). Blue light was delivered as soon as the assay was initiated at 7% O_2_. Locomotory speed responses to a switch from 7% O_2_ to 150 ppm H_2_S balanced with 7% O_2_ were recorded for day-one adults grown for 16 hours in the presence or absence of ATR. (**L**) Locomotory speed responses to a switch from 7% O_2_ to 150 ppm H_2_S balanced with 7% O_2_, following activation of O_2_ sensing neurons by expressing a gain-of-function allele of *pkc-*1 in O_2_ sensory neurons under *gcy-37* promoter. For the comparison of acute locomotory speed responses between strains or conditions, red bars on the x-axis represent two intervals (4–5 minutes and 11–12 minutes, indicated with red bars on the x-axis) for statistical analysis. **** = *p* < 0.0001, *** = *p* < 0.001, ** = *p* < 0.01, ns = not significant, Mann–Whitney U test.

**Figure 4—figure supplement 1.**
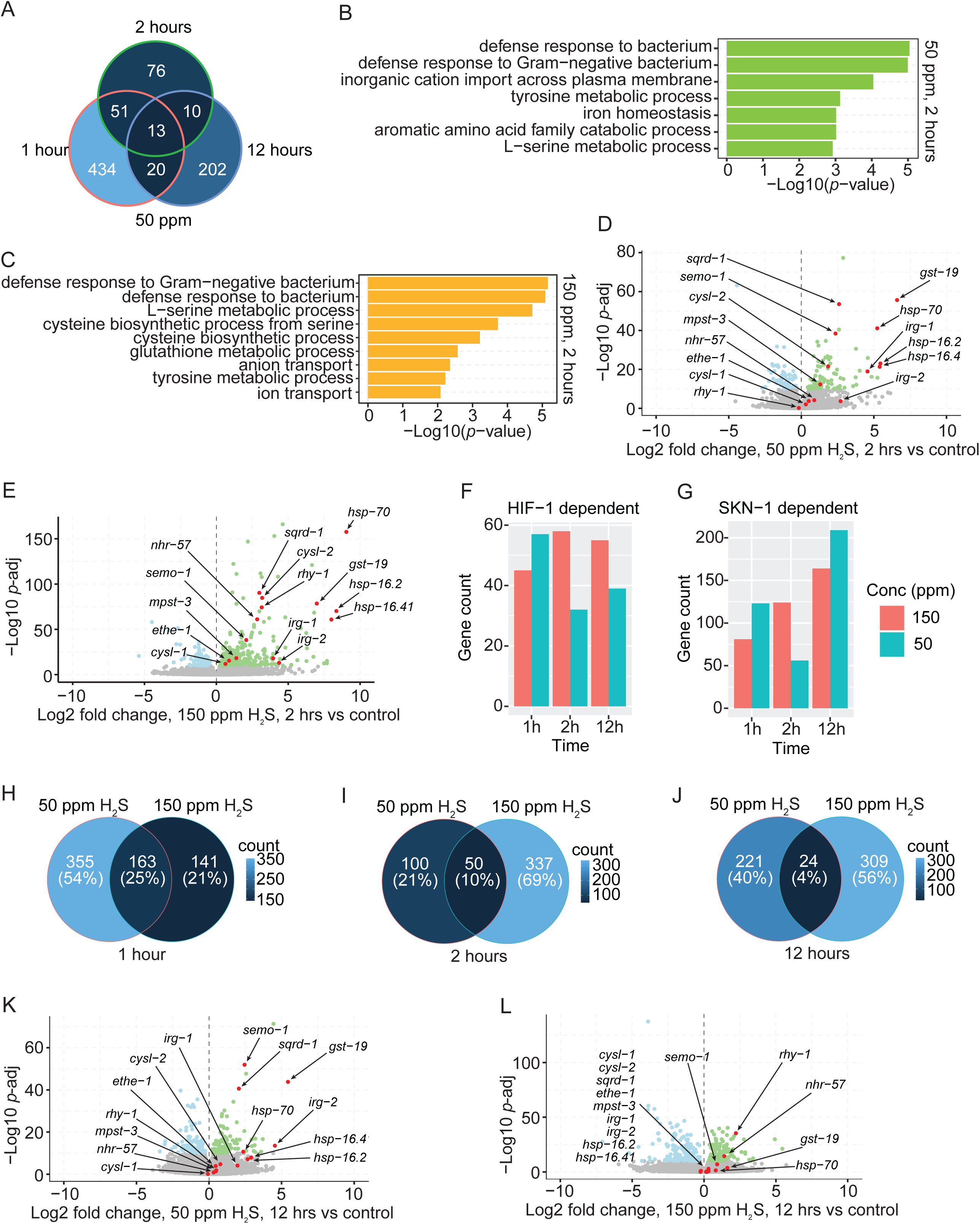
Transcriptome reprogramming induced by H_2_S exposure. (**A**) Venn diagram displaying the number of differentially expressed genes with adjusted *p* value <1e-10 in WT animals exposed to 50 ppm H_2_S balanced with 7% O_2_ for 1, 2, or 12 hours. (**B, C**) Significantly enriched GO categories for differentially expressed genes with adjusted *p* value <1e-10 in WT animals exposed to 50 ppm (**B**) or 150 ppm (**C**) H_2_S balanced with 7% O_2_ for 2 hours. (**D, E**) Volcano plots showing the differentially expressed genes with adjusted *p* value <1e-10 in WT animals exposed to 50 ppm (**D**) or 150 ppm (**E**) H_2_S balanced with 7% O_2_ for 2 hours. A set of genes involved in H_2_S detoxification, cysteine metabolism and stress response are highlighted in red. (**F, G**) The number of HIF-1(**F**) and SKN-1(**G**) target genes induced by 50 ppm or 150 ppm H_2_S exposure for 1 hour, 2 hours or 12 hours in WT animals, with adjusted *p* value < 1e-30. (**H–J**) Venn diagrams displaying the number of differentially expressed genes (adjusted *p* value <1e-10) in WT animals under the indicated conditions: animals exposed to 50 ppm or 150 ppm H_2_S for 1hour (**H**); 2 hours (**I**); and 12 hours (**J**). (**K, L**) Volcano plots showing the differentially expressed genes (adjusted *p* value <1e-10) in WT animals that were exposed to either 50 ppm (**K**) or 150 ppm (**L**) H_2_S balanced with 7% O_2_ for 12 hours. A set of genes involved in H_2_S detoxification, cysteine metabolism and stress response are highlighted in red.

**Figure 5—figure supplement 1.**
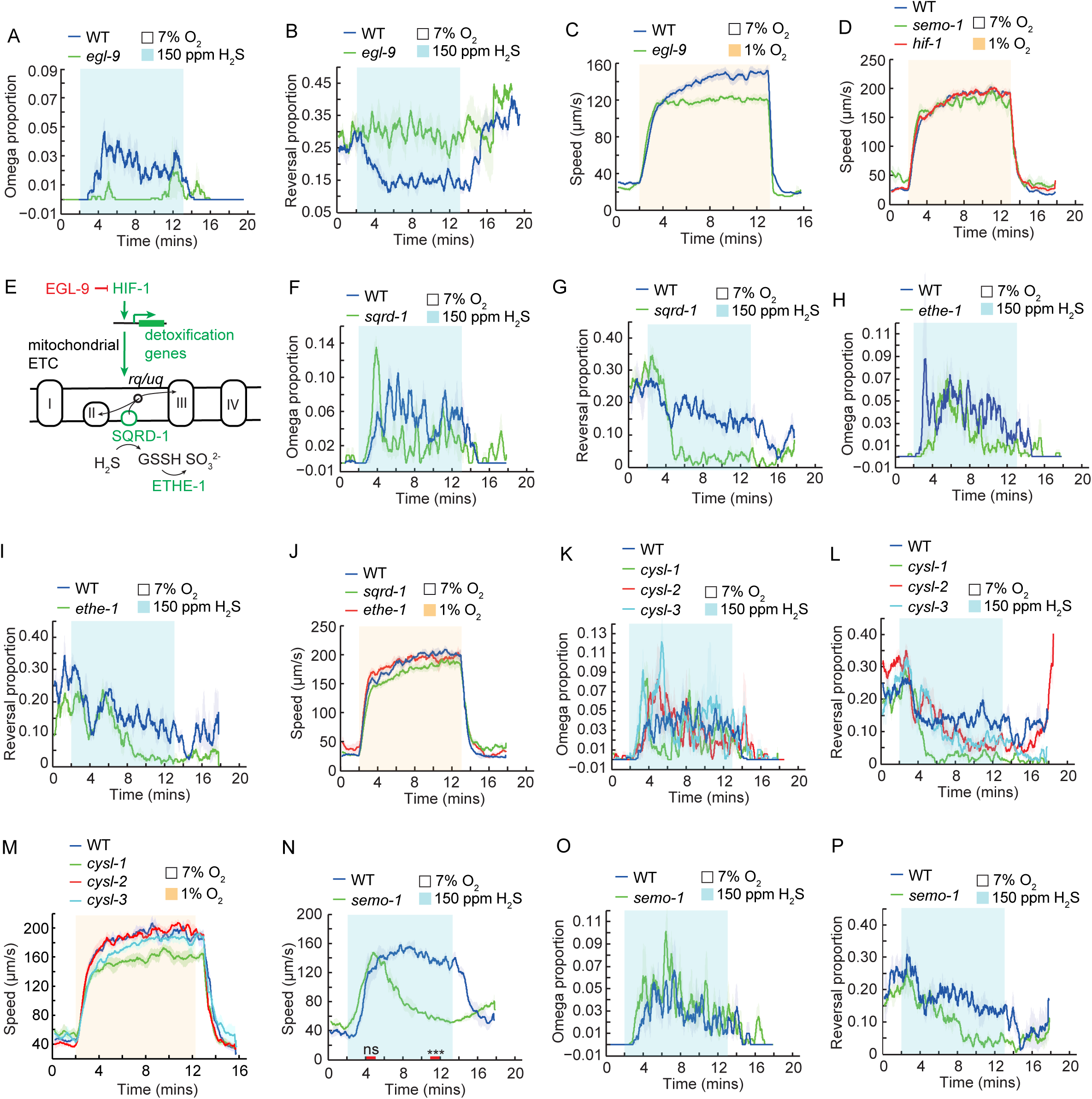
Acute response to H_2_S is modulated by HIF-1 signaling. (**A, B**) Fraction of WT and *egl-9(sa307)* animals undergoing reorientation movements (omega turns) (**A**) or backward locomotion (reversals) (**B**) during a switch from 7% O_2_ to 150 ppm H_2_S balanced with 7% O_2_. (**C, D**) Locomotory speed responses to a switch from 7% O_2_ to 1% O_2_ for WT and *egl-9(sa307)* (**C**); WT, *hif-1(ia4)*, and *semo-1(yum2889)* (**D**). (**E**) HIF-1 signaling regulates the H_2_S mitochondrial detoxification pathway. The detoxification enzyme SQRD-1 (sulfide:quinone oxidoreductase) oxidizes H_2_S, donating electrons to ubiquinone (uq), and rhodoquinone (rq) of the mitochondrial electron transport chain (ETC). ETHE-1 (persulfide dioxygenase) oxidizes sulfur compounds to sulfite. (**F–I**) Fraction of animals undergoing reorientation movements (omega turns, F and H) or backward locomotion (reversals, G and I) during a switch from 7% O_2_ to 150 ppm H_2_S balanced with 7% O_2_ for the indicated genotypes: WT and *sqrd-1(tm3378)* (**F, G**); and WT and *ethe-1(yum2895)* (**H, I**). (**J**) Locomotory speed responses to a switch from 7% O_2_ to 1% O_2_ for WT, *sqrd-1(tm3378)*, and *ethe-1(yum2895)*. (**K, L**) Fraction of WT, *cysl-1(ok762), cysl-2(ok3516)*, and *cysl-3(yum4)* animals undergoing reorientation movements (omega turns, K) or backward locomotion (reversals, L) during a switch from 7% O_2_ to 150 ppm H_2_S balanced with 7% O_2_. (**M**) Locomotory speed responses to a switch from 7% O_2_ to 1% O_2_ for WT, *cysl-1(ok762), cysl-2(ok3516)*, and *cysl-3(yum4)* animals. (**N**) Locomotory speed responses to a switch from 7% O_2_ to 150 ppm H_2_S balanced with 7% O_2_ for WT and *semo-1(yum2889).* (**O, P**) Fraction of animals undergoing reorientation (omega turns, **O**) or backward locomotion (reversals, **P**) in response to a switch from 7% O_2_ to 150 ppm H_2_S balanced with 7% O_2_. Comparisons of acute locomotory responses between strains were performed using two intervals (4–5 minutes and 11–12 minutes, indicated with red bars on the x-axis) for statistical analysis. *** = *p* < 0.001, ns = not significant, Mann–Whitney U test.

**Figure 6—figure supplement 1.**
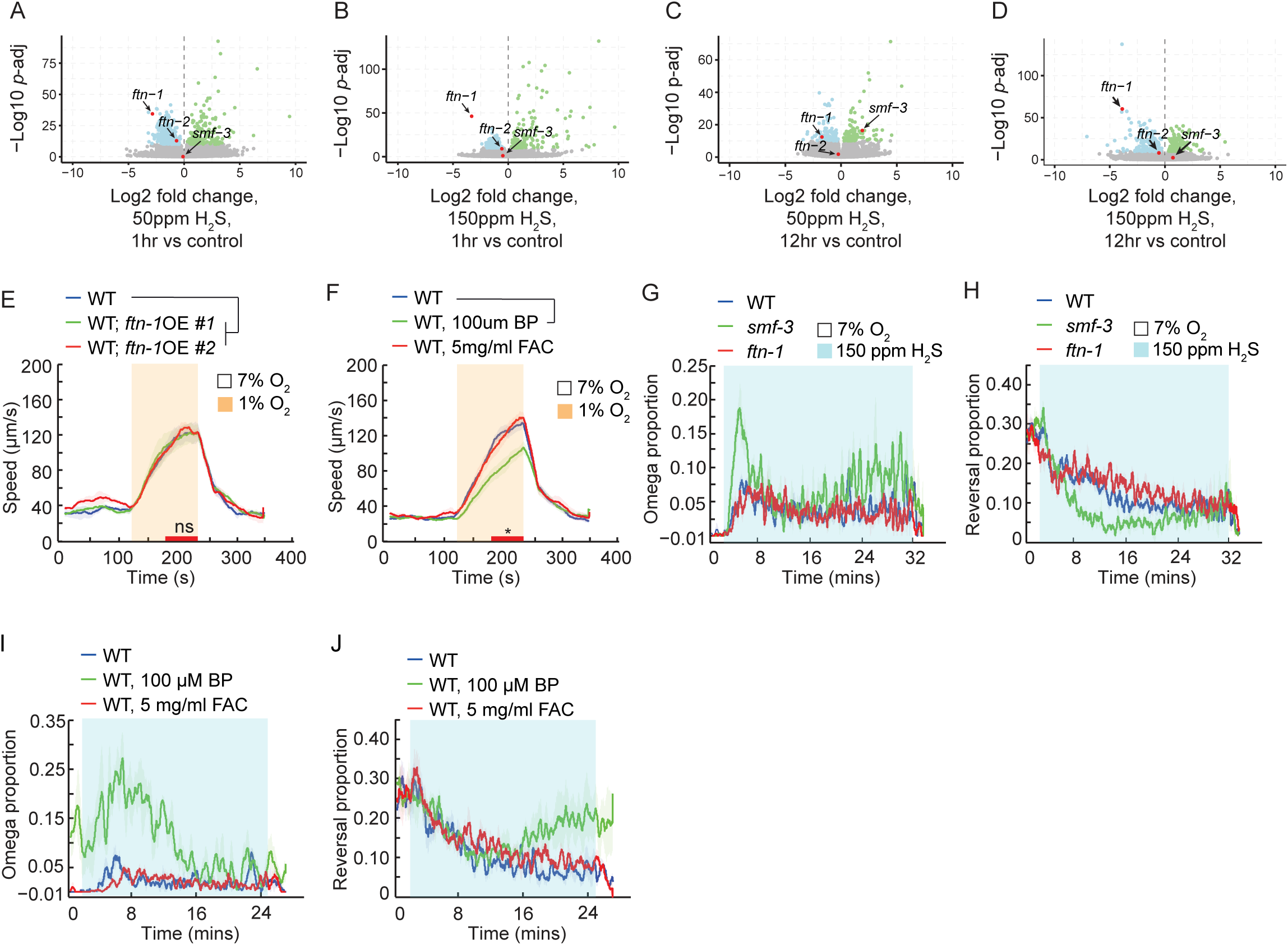
Labile iron pool sustains the locomotory activity in H_2_S. (**A–D**) Volcano plots showing the relative expression of the genes involved in the regulation of iron homeostasis (*ftn-1*, *ftn-2*, and *smf-3*) in WT animals under the following conditions: 1-hour exposure in 50 ppm H_2_S balanced with 7% O_2_ (**A**); 1-hour exposure in 150 ppm H_2_S balanced with 7% O_2_ (**B**); 12-hour exposure in 50 ppm H_2_S balanced with 7% O_2_ (**C**); and 12-hour exposure in 150 ppm H_2_S balanced with 7% O_2_ (**D**). (**E, F**) Locomotory speed responses to a switch from 7% O_2_ to 1% O_2_ for animals of the indicated genotype or treatment: WT, and two independent transgenic lines overexpressing *ftn-1* genomic DNA under its own promoter (#1 and #2) (**E**); WT animals, WT animals treated with 100 μM 2,2′-Bipyridyl (BP), and WT animals treated with 5 mg/ml ferric ammonium citrate (FAC) for 16 hours (**F**). Red bar on the x-axis represents one-minute time interval (3–4 minutes) used for statistical analysis. * = *p* < 0.05, ns = not significant, Mann–Whitney U test. (**G, H**) Fraction of animals undergoing reorientation movements (omega turns) (**G**) or backward locomotion (reversals) (**H**) during a switch from 7% O_2_ to 150 ppm H_2_S balanced with 7% O_2_ for animals of the indicated genotypes WT, *smf-3(ok1305)*, and *ftn-1(ok3625)*. (**I, J**) Fraction of animals undergoing reorientation movements (omega turns) (**I**) or backward locomotion (reversals) (**J**) during a switch from 7% O_2_ to 150 ppm H_2_S balanced with 7% O_2_ for WT animals, WT animals pretreated with either 100 μM 2,2′-Bipyridyl (BP) or with 5 mg/ml ferric ammonium citrate (FAC) in the presence of food for 16 hours.

**Figure 7—figure supplement 1.**
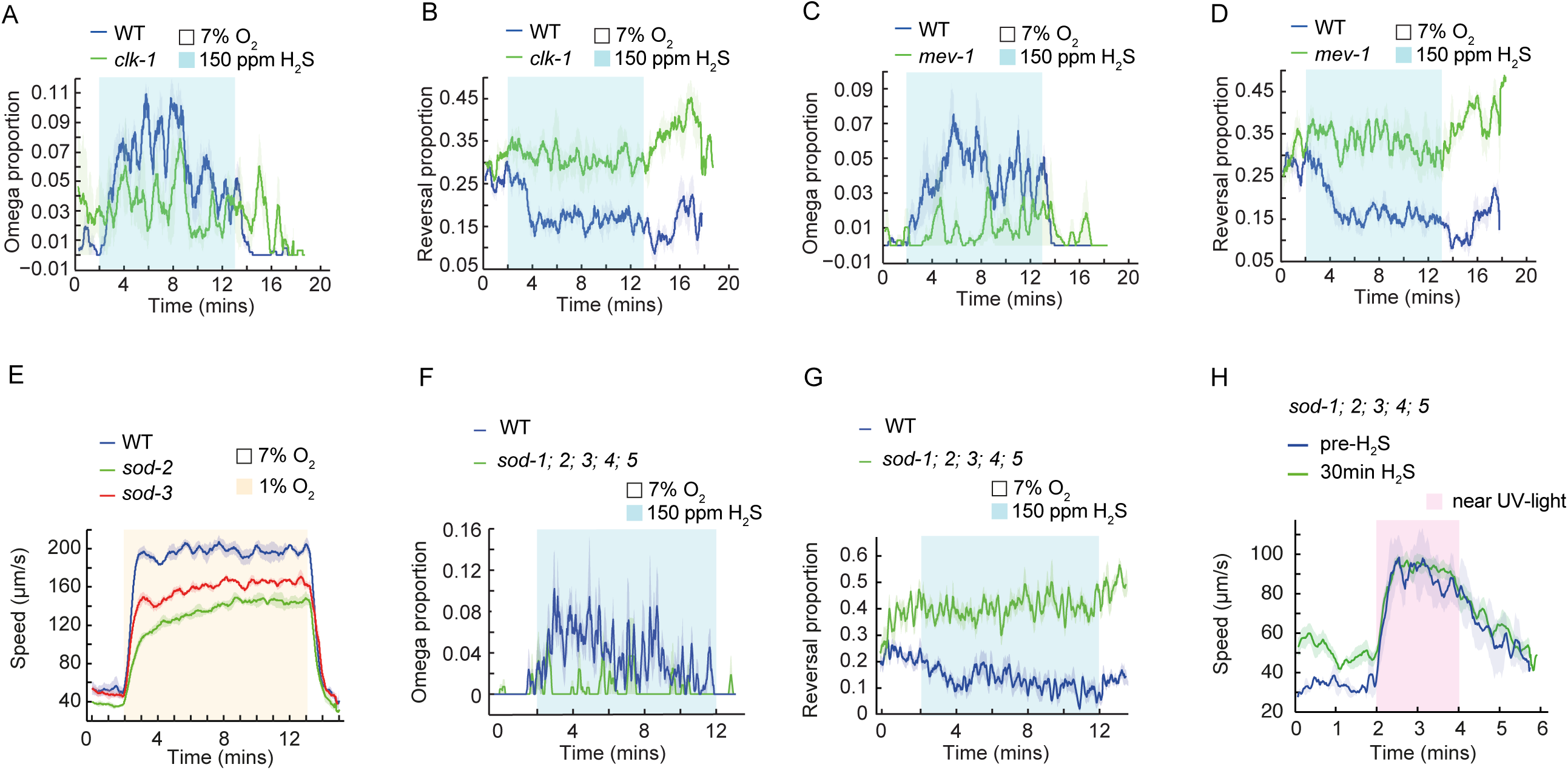
Mitochondrial function is required for acute response to H_2_S. (**A–D**) Fraction of animals undergoing reorientation movements (omega turns) or backward locomotion (reversals) during a switch from 7% O_2_ to 150 ppm H_2_S balanced with 7% O_2_ for the indicated genotype or treatment: WT and *clk-1(qm30)* (**A, B**); WT and *mev-1(kn1)* (**C, D**). (**E**) Locomotory speed responses to a switch from 7% O_2_ to 1% O_2_ for WT, *sod-2(ok1030);* and *sod-3(tm760)*. (**F, G**) Fraction of animals undergoing reorientation movements (omega turns, **F**) or backward locomotion (reversals, **G**) during a switch from 7% O_2_ to 150 ppm H_2_S balanced with 7% O_2_ for WT and *sod-1(tm783); sod-2(ok1030); sod-3(tm760); sod-4(gk101); sod-5 (tm1146*). (**H**) Locomotory response to acute near UV exposure for WT and *sod-1(tm783); sod-2(ok1030); sod-3(tm760); sod-4(gk101); sod-5 (tm1146*) before and after 30 minutes of exposure to 150ppm H_2_S.

**Figure 1—Video 1. The locomotory response of WT animals to H_2_S**

An 8x accelerated video clip (original length: 16 minutes) showing WT (N2) animals exposed to 7% O_2_ for 2 minutes, followed by 11 minutes in 150 ppm H_2_S, and then returned to 7% O_2_. The H_2_S exposure segment is highlighted and begins at the 15-second mark, corresponding to 2 minutes in the original recording.

