## Supplementary file 1 for "Avoidance of hydrogen sulfide is modulated by external and internal states in *C. elegans*"

| Groups | Strain name | Genes | Locomotory response | Omega  Proportion | Reversal  Proportion |
| --- | --- | --- | --- | --- | --- |
| Guanylate cyclase |  | *gcy-1(tm2669)* | +++ | +++ | +++ |
|  | VC3024 | *gcy-2(ok3721)* | +++ | +++ | +++ |
|  | VC2796 | *gcy-3(gk1154)* | +++ | +++ | +++ |
|  |  | *gcy-4(tm1653)* | +++ | +++ | +++ |
|  | RB1010 | *gcy-5(ok930)* | +++ | +++ | +++ |
|  | OH4844 | *gcy-5(tm897)* | +++ | +++ | +++ |
|  |  | *gcy-6(tm1449)* | ++ | +++ | +++ |
|  |  | *gcy-7(tm901)* | +++ | +++ | +++ |
|  | IK800 | *gcy-8(oy44)* | +++ | +++ | +++ |
|  |  | *gcy-9(tm7632)* | +++ | +++ | +++ |
|  |  | *gcy-11(tm8150)* | +++ | +++ | +++ |
|  | CHS2307 | *gcy-12(yum2919)* | +++ | +++ | +++ |
|  | CHS1723 | *gcy-13(yum2920)* | +++ | +++ | +++ |
|  | JN1194 | *gcy-14(pe1102)* | +++ | +++ | +++ |
|  |  | *gcy-14(tm4210)* | +++ | +++ | +++ |
|  |  | *gcy-14(tm12274)* | +++ | +++ | +++ |
|  | VC2675 | *gcy-15(gk1102)* | +++ | +++ | +++ |
|  | VC2450 | *gcy-17(gk1155)* | +++ | +++ | +++ |
|  |  | *gcy-17(tm4516)* | +++ | +++ | +++ |
|  | IK597 | *gcy-18(nj38); gcy-8(oy44); gcy-23(nj37)* | +++ | +++ | +++ |
|  | RB1909 | *gcy-19(ok2472)* | +++ | +++ | +++ |
|  | RB1935 | *gcy-20(ok2538)* | ++ | ++ | +++ |
|  | CHS2335 | *gcy-20(yum2921)* | +++ | +++ | +++ |
|  |  | *gcy-21(tm11147)* | +++ | +++ | +++ |
|  | CHS2325 | *gcy-21(yum2922)* | +++ | +++ | +++ |
|  |  | *gcy-22(tm2364)* | ++ | +++ | +++ |
|  | RB924 | *gcy-23(ok797)* | ++ | +++ | +++ |
|  | IK427 | *gcy-23(nj37)* | +++ | +++ | +++ |
|  |  | *gcy-25(tm4300)* | +++ | +++ | +++ |
|  | CHS2336 | *gcy-27(yum2923)* | +++ | +++ | +++ |
|  | CHS502 | *gcy-28(yum32)* | +++ | +++ | +++ |
|  | CHS2326 | *gcy-29(yum2924)* | +++ | +++ | +++ |
|  | DR47 | *daf-11(m84)* | - | ++ | - |
|  | CX2065 | *odr-1(n1963)* | +++ | +++ | +++ |
|  | CZ3714 | *gcy-31(ok296)* | +++ | +++ | +++ |
|  | RB1048 | *gcy-32(ok995)* | +++ | +++ | +++ |
|  | CZ3715 | *gcy-33(ok232)* | ++ | +++ | +++ |
|  | RB1062 | *gcy-34(ok1012)* | +++ | +++ | +++ |
|  | AX1295 | *gcy-35(ok769)* | +++ | +++ | +++ |
|  | [AX1296](https://cgc.umn.edu/strain/AX1296) | *gcy-36(db42)* | +++ | +++ | +++ |
|  | RB626 | *gcy-37(ok384)* | +++ | +++ | +++ |
| Globins | CHS506 | *glb-1(yum12)* | +++ | +++ | +++ |
|  | CHS507 | *glb-2(yum13)* | +++ | +++ | +++ |
|  | CHS539 | *glb-3(yum29)* | +++ | +++ | +++ |
|  | CHS525 | *glb-4(yum22)* | +++ | +++ | +++ |
|  |  | *glb-5(tm5440)* | +++ | +++ | +++ |
|  |  | *glb-6(tm3795)* | +++ | +++ | +++ |
|  | CHS535 | *glb-7(yum27)* | +++ | +++ | +++ |
|  | CHS541 | *glb-8(yum30)* | +++ | +++ | +++ |
|  | CHS519 | *glb-9(yum19)* | +++ | +++ | +++ |
|  |  | *glb-10(tm5198)* | +++ | +++ | +++ |
|  |  | *glb-10(tm5744)* | +++ | +++ | +++ |
|  | CHS509 | *glb-11(yum14)* | ++ | +++ | +++ |
|  | CHS543 | *glb-12(yum31)* | +++ | +++ | +++ |
|  |  | *glb-13(tm2825)* | +++ | +++ | +++ |
|  | CHS2204 | *glb-14(yum594)* | +++ | +++ | +++ |
|  | CHS511 | *glb-15(yum15)* | +++ | +++ | +++ |
|  |  | *glb-16(tm5264)* | +++ | +++ | +++ |
|  | CHS513 | *glb-17(yum16)* | +++ | +++ | +++ |
|  |  | *glb-18(tm5798)* | +++ | +++ | +++ |
|  |  | *glb-18(tm6017)* | +++ | +++ | +++ |
|  |  | *glb-19(tm6965)* | +++ | +++ | +++ |
|  |  | *glb-20(tm2286)* | +++ | +++ | +++ |
|  |  | *glb-21(tm8033)* | +++ | +++ | +++ |
|  | CHS529 | *glb-22(yum24)* | +++ | +++ | +++ |
|  | CHS515 | *glb-23(yum17)* | +++ | +++ | +++ |
|  | CHS517 | *glb-24(yum18)* | +++ | +++ | +++ |
|  | CHS531 | *glb-25(yum25)* | +++ | +++ | +++ |
|  |  | *glb-26(tm4837)* | +++ | +++ | +++ |
|  | CHS521 | *glb-27(yum20)* | +++ | +++ | +++ |
|  |  | *glb-28(tm6910)* | +++ | +++ | +++ |
|  | CHS527 | *glb-29(yum23)* | +++ | +++ | +++ |
|  | CHS537 | *glb-30(yum28)* | ++ | +++ | +++ |
|  | CHS523 | *glb-31(yum21)* | +++ | +++ | +++ |
|  | CHS533 | *glb-32(yum26)* | +++ | +++ | +++ |
|  |  | *glb-33(tm3656)* | +++ | +++ | +++ |
| Potassium channels | CHS5086 | *shk-1; shl-1* | ++ | +++ | +++ |
|  | CHS5067 | *exp-2; kvs-1; kvs-2; kvs-3; kvs-4; kvs-5* | ++ | +++ | +++ |
|  | CHS5055 | *shw-1; egl-36; shw-3* | +++ | +++ | +++ |
|  | CHS5047 | *kqt-1; kqt-2; kqt-3* | +++ | +++ | +++ |
|  | CHS5073 | *egl-2; unc-103* | +++ | +++ | +++ |
|  | CHS5100 | *slo-1; slo-2* | +++ | +++ | +++ |
|  | CHS5057 | *kcnl-1; kcnl-2; kcnl-3; kcnl-4* | +++ | +++ | +++ |
|  | CHS5115 | *egl-23; twk-9* | +++ | +++ | +++ |
|  | CHS5105 | *sup-9; twk-20* | +++ | +++ | +++ |
|  | CHS5069 | *unc-58; unc-110* | ++ | +++ | +++ |
|  | CHS5100 | *twk-1; twk-2* | ++ | +++ | +++** |
|  | CHS5121 | *twk-3; twk-10* | +++ | +++ | +++ |
|  | CHS5061 | *twk-4; twk-5* | +++ | +++ | +++ |
|  | CHS5085 | *twk-6* | +++ | +++ | +++ |
|  | CHS5049 | *twk-7; twk-8; twk-40* | + | +++ | +++ |
|  | CHS5053 | *twk-11; twk-12; twk-13* | +++ | +++ | +++ |
|  | CHS5014 | *twk-14; twk-16; twk-17* | +++ | +++ | +++ |
|  | CHS5031 | *twk-21; twk-22; twk-23* | +++ | +++ | +++ |
|  | CHS5006 | *twk-24; twk-26; twk-45* | +++ | +++ | +++ |
|  | CHS5043 | *twk-25; twk-33; twk-34; twk-36* | +++ | +++ | +++ |
|  | CHS5038 | *twk-28; twk-29; twk-30* | +++ | +++ | +++ |
|  | CHS5025 | *twk-31; twk-32; twk-35* | +++ | +++ | +++ |
|  | CHS5081 | *twk-37; twk-39; twk-48* | + | +++ | +++ |
|  | CHS5045 | *twk-42; twk-43; twk-44* | +++ | +++ | +++ |
|  | CHS5019 | *twk-46; twk-47; twk-49* | +++ | +++ | +++ |
|  |  | *irk-1; irk-2; irk-3* | +++ | +++ | +++ |
| Ion channels | PR691 | *tax-2(p691)* | + | +++ | +++ |
|  | PR694 | *tax-2(p694)* | + | ++ | +++ |
|  | PR678 | *tax-4(p678)* | + | +++ | +++ |
|  | CHS1551 | *cng-1(yum1117)* | +++ | +++ | +++ |
|  | CHS1552 | *cng-2(yum1118)* | +++ | +++ | +++ |
|  | CHS1553 | *cng-3(yum1119)* | +++ | +++ | +++ |
|  | CB1126 | *cng-4(e1126)* | +++ | +++ | +++ |
|  |  | *cng-4(tm5036)* | +++ | +++ | +++ |
|  | KJ5560 | *cng-1(jh111); cng-3(jh113)* | +++ | +++ | +++ |
|  | KJ5560 | *cng-1(jh111); cng-3(jh113); tax-4(p678)* | + | +++ | +++ |
|  | [VC1233](https://cgc.umn.edu/strain/VC1233) | *ocr-2(ok1711)* | +++ | +++ | +++ |
|  | [CX4544](https://cgc.umn.edu/strain/CX4544) | *ocr-2(ak47)* | +++ | +++ | +++ |
| Neuropeptide processing enzymes | AX2210 | *egl-21(n476)* | - | +++ | +++ |
|  | VC671 | *egl-3(ok979)* | + | - | +++ |
| Cilia mutants | OE3063 | *daf-19(m86)* | - | ++ | -* |
|  | SP1603 | *dyf-3(m185)* | + | ++ | -* |
|  | SP1196 | *dyf-7(m539)* | - | + | +++* |
| Hypoxia-pathway related mutants | ZG31 | *hif-1(ia4)* | ++ | +++ | +++** |
|  | JT307 | *egl-9(sa307)* | - | - | - |
|  | CB5602 | *vhl-1(ok161)* | - | +++ | - |
|  | CB6088 | *egl-9(sa307); hif-1(ia4)* | ++ | ++ | +++** |
|  | CHS340 | *vhl-1(ok161); hif-1(ia4)* | ++ | +++ | +++** |
| Mitochondrial related mutants | CW152 | *gas-1(fc21)* | - | + | ++ |
|  | MQ130 | *clk-1(qm30)* | - | ++ | -* |
|  | TK22 | *mev-1(kn-1)* | - | + | -* |
|  | CHS2184 | *isp-1(qm150)* | - | - | -* |
| Biogenic amines | CHS2309 | *tph-1(yum99)* | +++ | +++ | +++ |
|  | GRB21 | *tph-1(mg280)* | +++ | +++ | +++ |
|  |  | *cat-2(tm2261)* | +++ | +++ | +++ |
|  | RB993 | *tdc-1(ok914)* | +++ | +++ | +++ |
|  | RB1161 | *tbh-1(ok1196)* | +++ | +++ | +++ |
|  | MT9455 | *tbh-1(n3247)* | +++ | +++ | +++ |
| Neurotransmisson  related mutants | MT6308 | *eat-4(ky5)* | ++ | +++ | +++ |
|  | KP4 | *glr-1(n2461)* | +++ | +++ | +++ |
|  | [RM2710](https://cgc.umn.edu/strain/RM2710) | *snf-11(ok156)* | +++ | +++ | +++ |
|  | PR1152 | *cha-1(p1152)* | - | +++ | +++ |
|  | VC862 | *cho-1(ok1069)* | + | +++ | +++ |
| Superoxide dismutase | FX776 | *sod-1(tm776)* | +++ | +++ | +++ |
|  | RB1072 | *sod-2* *(ok1030)* | + | +++ | +++ |
|  | GA186 | *sod-3(tm760)* | + | +++ | +++ |
|  | GA416 | *sod-4(gk101)* | +++ | +++ | +++ |
|  | GA503 | *sod-5(tm1146)* | ++ | +++ | +++ |
|  | MQ1766 | *sod-1; sod-2; sod-3; sod-4; sod-5* | - | + | -** |
| Wild isolates | PX179 |  | ++ | ++ | ++ |
|  | PX174 |  | + | + | - |
|  | PB303 |  | + | - | -* |
|  | ED3077 |  | + | - | + |
|  | ED3014 |  | + | - | - |
|  | ED3017 |  | + | - | - |
|  | EG4946 |  | ++ | ++ | ++ |
|  | LKC34 |  | +++ | +++ | +++ |
|  | JU1088 |  | +++ | +++ | +++ |
|  | JU1543 |  | ++ | + | ++ |
|  | JU258 |  | +++ | +++ | +++ |
|  | JU561 |  | + | - | - |
|  | MY10 |  | ++ | + | +++* |
|  | MY16 |  | + | - | -* |
|  | KY314 |  | +++ | +++ | +++ |
|  | CB4856 |  | ++ | +++ | +++* |
|  | CB4858 |  | - | + | -* |
|  | CB4855 |  | + | ++ | +++* |
| Others | CB1370 | *daf-2(e1370)* | - | - | -* |
|  | GR1352 | *daf-16(mgDf47)* | +++ | +++ | +++ |
|  | GR1309 | *daf-2(e1370); daf-16(mgDf47)* | +++ | +++ | +++ |
|  | GR1311 | *daf-3(mgDf90)* | +++ | +++ | +++ |
|  | CB1372 | *daf-7(e1372)* | - | + | - |
|  |  | *daf-3(mgDf90); daf-7(e1372)* | +++ | +++ | +++ |
|  | AX204 | *npr-1(ad609)* | - | + | ++ |
|  | AX1197 | *npr-1(ad609); gcy-35(ok769)* | +++ | +++ | +++ |
|  | AX625 | *npr-1(ad609); tax-4(p678)* | + | ++ | +++ |
|  | RB1667 | *tax-6(ok2065)* | - | +++ | + |
|  | VC990 | *cnb-1(ok276)* | - | + | ++ |
|  | STE68 | *nhr-49(nr2041)* | + | +++ | +++ |
|  |  | *sqrd-1(tm3378)* | + | ++ | +++** |
|  | CHS1657 | *ethe-1(yum2895)* | +++ | ++ | +++** |
|  | CHS1635 | *semo-1(yum2889)* | +++ | +++ | +++** |
|  | RB1074 | *smf-3(ok1305)* | +++ | +++ | +++** |
|  | RB2603 | *ftn-1(ok3625)* | ++ | +++ | +++ |
|  | CHS1726 | *hif-1(ia4); ftn-1(yum2918)* | +++ | ++ | +++** |
|  | PR680 | *che-1(p680)* | +++ | ++ | +++ |
|  | CB1033 | *che-2(e1033)* | + | +++ | ++ |
|  | PR802 | *osm-3(p802)* | +++ | +++ | +++ |
|  | CX4 | *odr-7(ky4)* | +++ | +++ | +++ |
|  | CHS395 | *cbs-1(yum10)* | + | + | + |
|  | RB839 | *cbs-2(ok666)* | +++ | +++ | +++ |
|  | VC2569 | *cth-1(ok3319)* | +++ | +++ | +++ |
|  | GR2257 | *cth-2(mg599)* | +++ | +++ | +++ |
|  | CHS1722 | *cysl-1(ok762)* | + | +++ | +++** |
|  | RB2535 | *cysl-2(ok3516)* | + | +++ | +++** |
|  | CHS345 | *cysl-3(yum4)* | + | +++ | +++** |
|  | RB2436 | *cysl-4(ok3359)* | ++ | +++ | +++ |
|  | CHS346 | *mpst-2(yum5)* | +++ | +++ | +++ |
|  |  | *mpst-3(tm4387)* | ++ | +++ | +++ |
|  | CHS347 | *mpst-4(yum6)* | +++ | +++ | +++ |

- 0 to 10% of wild type response to 150ppm H_2_S in 7% O_2_

+	10 to 50% of wild type response to 150ppm H_2_S in 7% O_2_

++	50 to 80% of wild type response to 150ppm H_2_S in 7% O_2_

+++ 80 to 100% of wild type response or higher response to 150ppm H_2_S in 7% O_2_

*	Mutants with higher basal reversal

**	Mutants with a more rapid decline in reversal

Mutants with an enhanced initial omega-turn response
